# *FOXP2* confers oncogenic effects in prostate cancer through activating MET signalling

**DOI:** 10.1101/2022.07.06.498943

**Authors:** Xiaoquan Zhu, Chao Chen, Dong Wei, Yong Xu, Siying Liang, Wenlong Jia, Jian Li, Yanchun Qu, Jianpo Zhai, Yaoguang Zhang, Pengjie Wu, Qiang Hao, Linlin Zhang, Wei Zhang, Xinyu Yang, Lin Pan, Ruomei Qi, Yao Li, Feiliang Wang, Rui Yi, Ze Yang, Jianye Wang, Yanyang Zhao

## Abstract

Identification oncogenes is fundamental to revealing the molecular basis of cancer. Here, we found that *FOXP2* is overexpressed in human prostate cancer cells and prostate tumors, but its expression is absent in normal prostate epithelial cells and low in benign prostatic hyperplasia. To date, little is known regarding the link of *FOXP2* to prostate cancer. We observed that high *FOXP2* expression and frequent amplification are significantly associated with high Gleason score. Ectopic expression of *FOXP2* induces malignant transformation of mouse NIH3T3 fibroblasts and human prostate epithelial cell RWPE-1. Conversely, *FOXP2* knockdown suppresses the proliferation of prostate cancer cells. Transgenic overexpression of *FOXP2* in the mouse prostate causes prostatic intraepithelial neoplasia. Overexpression of *FOXP2* aberrantly activates oncogenic MET signalling and inhibitors targeting MET signalling effectively reverts the *FOXP2*-induced oncogenic phenotype. Additionally, the novel recurrent *FOXP2-CPED1* fusion identified in prostate tumors results in high expression of truncated FOXP2, which exhibit a similar capacity for malignant transformation. Together, our data demonstrate for the first time that *FOXP2* is an oncogene involved in tumorigenicity of prostate.

## Introduction

Oncogenes arise as a consequence of genetic alterations that increase the expression level or activity of a proto-oncogene. Activation of oncogenes causes malignant transformation of normal cells and maintains cancerous cell proliferation and survival. Clinical intervention strategies targeting oncoproteins and their related signalling pathways lead to growth arrest and apoptosis in cancer cells [1, 2].

Prostate cancer is the second most commonly diagnosed cancer worldwide in men, with ∼1.3 million new cases and ∼ 0.36 million deaths in 2018 [3]. In China, the incidence rate of prostate cancer has increased quickly, with an annual percentage change of ∼ 13% since 2000 [4]. These findings stress the importance of a comprehensive understanding of the molecular mechanisms underlying the genesis of prostate cancer. Accumulating data from genomic characterization and functional studies have revealed some oncogenes, including the E26 transformation-specific (ETS) family genes *ERG* and *ETV1*, among others, whose overexpression is involved in prostate tumorigenesis [5–7]. However, the oncogenes that contribute to the initiation of the disease remain unclear.

In this study, we identified a previously unknown intrachromosomal gene fusion involving *FOXP2* in primary prostate tumors by RNA-Seq and RT-PCR combined with sanger-sequencing. The fusion encodes a truncated FOXP2 mutant protein encompassing the key domains and is highly expressed in the fusion-carrying prostate tumor. *FOXP2* is tightly associated with vocal development in humans and mammals [8–10]. Mutations in *FOXP2* are the only known cause of developmental speech and language disorder in humans [11]. Most of the recent studies investigating the roles of *FOXP2* in cancer have focused on noncoding RNA-mediated dysregulation of *FOXP2*, suggesting that *FOXP2* has either oncogenic or tumor-suppressive effects on cancer development in different tumor contexts [12–16]. However, to date, little is known regarding the link between *FOXP2* and prostate cancer. Here, we investigated the expression of *FOXP2* in prostate cancer samples. Moreover, we explored the biological functions of *FOXP2* in human prostate tumors and prostate cancer cell lines. In addition, the impact of prostate-specific expression of the *FOXP2* gene in mice was tested.

## Materials and Methods

### Clinical Specimen

One hundred localized primary prostatic adenocarcinomas that underwent radical prostatectomy were collected. All tumor specimens were independently evaluated by two pathologists for histological diagnosis and Gleason score on haematoxylin and eosin (H&E) stained slides. The presence of or morphological absence of adenocarcinoma was determined by one pathologist through review of the 4 μm H&E stained tissue slice, and then high-density cancer foci (more than 70% tumor cellularity) and contamination-free normal prostate tissues were cut into ∼1 mm^3^ tissue blocks for extraction of DNA, RNA or protein. Ten pairs of prostate tumors (PC_1 to PC_10) and matched normal tissues (NT_1 to NT_10) were used for transcriptome sequencing (Table S1). Additionally, whole-genome sequencing was performed on the *FOXP2*-*CPED1* fusion-positive tumor (PC_1). Reduced Representation Bisulfite Sequencing (RRBS) analysis and small RNA sequencing were performed on PC_1 and NT_1.

### Cell Lines and Cell Culture

NIH3T3 mouse primary fibroblast cells, the immortalized human prostate epithelial cell line RWPE-1, prostate cancer cell line LNCaP and PC3 were obtained from the Cell Resource Center, Peking Union Medical College, National Infrastructure of Cell Line Resource. Prostate cancer cell line VCaP and Human embryonic kidney (HEK) 293T cells were obtained from ATCC. The identity of the cell lines was authenticated with STR profiling. The cell lines were checked free of mycoplasma contamination by PCR. LNCaP cells and HEK 293T cells were cultured in RPMI-1640 (Life Technologies) supplemented with 10% FBS (Life Technologies), L-glutamine (2 mM) (Life Technologies), penicillin (100 U/ml) (Life Technologies), and streptomycin (100 μg/ml) (Life Technologies). PC3 cells were cultured in F-12K (Life Technologies) supplemented with 10% FBS (Life Technologies), L-glutamine (2 mM) (Life Technologies), penicillin (100 U/ml) (Life Technologies), and streptomycin (100 μg /ml) (Life Technologies). NIH3T3 cells and VCaP cells were maintained in DMEM (Life Technologies) supplemented with 10% FBS, L-glutamine (2 mM), penicillin (100 U/ml), and streptomycin (100 μg /ml). RWPE-1 cells were grown in Defined Keratinocyte-serum free medium (Defined Keratinocyte-SFM, Life Technologies) with L-glutamine (2 mM), penicillin (100 U/ml), and streptomycin (100 μg /ml). All cells were grown at 37 °C.

### RNA Sequencing of Localized Prostate Cancer Samples

Total RNAs of 10 pairs of tumors (PC_1 to PC_10) and matched normal tissues (NT_1 to NT_10) were extracted with Trizol (Life Technologies). RNA integrity number (RIN) > 7.0 and a 28S: 18S ratio > 1.8. Sequencing libraries for strand-specific transcriptome was carried out as described previously [17] by BGI-Shenzhen (Shenzhen, China). Briefly, Beads containing oligo (dT) were used to isolate poly (A) mRNA from total RNA. The mRNA was fragmented into short fragments by the fragmentation buffer. Using these short fragments as templates, random hexamer-primers were used for synthesization of the first-strand cDNA. After purification with the G-50 gel filtration spin-column (GE Healthcare Life Sciences) to remove dNTPs, second-strand synthesis was performed by incubation with RNase H, DNA polymerase, and dNTPs containing dUTP (Promega). A single 3’ “A” base was added to the end-repaired cDNA. Upon ligation with the Illumina PE adaptors, the products were gel-recovered and subsequently digested with N-Glycosylase (UNG; Applied Biosystems) to remove the second-strand cDNA. Samples were then amplified with Phusion polymerase and PCR primers of barcode sequence. The amplified library was sequenced on an Illumina HiSeq 2000 sequencing platform. The paired-end reads obtained from HiSeq 2000 were aligned to the human reference genome and transcriptome (hg19) using SOAP2 program [18]. No more than 5 mismatches were allowed in the alignment for each read. The gene expression level was calculated by using RPKM [19] (Reads per kilobase transcriptome per million mapped reads), and the formula was shown as follows:

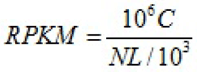

Given to be the expression of gene A, C represents number of reads that are uniquely aligned to gene A, N represents total number of reads that are uniquely aligned to all genes, and L represents number of bases on gene A. Using “The significance of digital gene expression profiles” [20], we identified differentially expressed genes between the tumors and matched normal samples based on the following criteria: FDR ≤ 0.001 and fold change ≥ 1.5.

### Detecting Human Fusion Genes in Localized Prostate Cancer Samples

We used SOAPfuse [21] to detect gene fusion events from RNA-Seq data based on the default parameters. After mapping RNA-Seq reads to the human reference genome sequence (hg19) and Ensembl annotated genes (release 64), SOAPfuse sought span-reads and junc-reads to support fusion detection: paired-end reads that mapped to two different genes were defined as span-reads, and reads covering the junction sites were called as junc-reads. SOAPfuse detects fusion events generated by genome rearrangements with breakpoints in intron and exon regions.

### Small RNA Sequencing of Localized Prostate Cancer Sample

Total RNAs of the tumor (PC_1) and matched normal tissue (NT_1) were extracted with Trizol (Life Technologies). Preparation of small RNA library was performed according to the manufacturer’s instructions (Preparing Samples for Analysis of Small RNA, Part #11251913, Rev. A) by BGI-Shenzhen (Shenzhen, China). Briefly, small RNA sequence ranging from 18 to 30 nt was gel-purified and ligated to the Illumina 3’ adaptor and 5’ adaptor. Ligation products were then gel-purified, reverse transcribed, and amplified. The amplified library was sequenced on an Illumina HiSeq 2000 platform. The small RNA reads were subjected to the following filtering processes: (i) Filtering out low-quality reads; (ii) Trimming 3’ adaptor sequence; (iii) Removing adaptor contaminations resulted from adaptor ligation; (iv) Retaining only short trimmed reads of sizes from 18 to 30 nt. To annotate and categorize small RNAs into different classes, we filtered out small RNA reads which might be from known noncoding RNAs by comparing them with known noncoding RNAs, including rRNA, tRNA, snRNA and snoRNA, which were deposited in the Rfam database and the NCBI Genbank. Small RNA reads assigned to exonic regions were also discarded. After removing small RNA reads in term of the above categories, the rest were subjected to MIREAP, which identified miRNA candidates according to the canonical hairpin structure and sequencing data. The identified small RNA (miRNA) reads were then aligned to miRNA reference sequences with tolerance for one mismatch. Reads that were uniquely aligned and overlapped with known miRNAs were considered as candidate miRNAs.

### Whole-Genome Sequencing of Primary Prostate Cancer Sample

Genomic DNA of PC_1 was extracted with phenol-chloroform method and subjected to whole-genome sequencing by BGI-Shenzhen (Shenzhen, China). After removing reads that contained sequencing adapters and low-quality reads with more than five ambiguous bases, high-quality reads were aligned to the NCBI human reference genome (hg19) using BWA (v0.5.9) [22] with default parameters. Picard (v1.54) was used to mark duplicates and followed by Genome Analysis Tool kit (v1.0.6076, GATK IndelRealigner) [23] was followed to improve alignment accuracy. The SNVs were detected by SOAP snp [24] and several filtering steps were performed to reduce the false positives, including the removal of SNVs whose consensus quality was lower than 20, single nucleotide variations (SNVs) located within 5 bp of the splice donor sites, and SNVs with less than 3 spanning reads. We next used BreakDancer to detect structural variations (SVs). After SVs were identified, we used ANNOVAR to do annotation and classification.

### Reduced Representation Bisulfite Sequencing of Primary Prostate Cancer Sample

Genomic DNAs of PC_1 and NT_1 were subjected to RRBS by BGI-Shenzhen (Shenzhen, China). Before library construction, MspI (NEB R0106L) was used to digest the genomic DNA. Next, the Illumina Paired-End protocol was used to construct the library. DNA libraries of 40-220 bp were excised. Then the excised DNA was recovered by columns, purified by MiniElute PCR Purification Kit (QIAGEN, #28006), and eluted in EB. Bisulfite was converted using the EZ DNA Methylation-Gold kit (ZYMO). All the bisulfite-converted products were amplified by PCR in a final reaction. PCR products were purified and recovered, followed by sequencing with an Illumina HiSeq 2000 platform. The RRBS reads were aligned to human genome reference (hg19) using SOAP2 allowing less than two mismatches. We then used these uniquely mapped reads that contained the enzyme cutting site to get methylation information of cytosine as described previously [25]. Methylation level was determined by dividing the number of reads covering each methyl-cytosine by the total reads which covered cytosine, and the methylation information for each gene were calculated when the promoters of which were covered by at least 5 CpGs.

### RT**-**PCR, qPCR, 3’RACE**-**PCR and Long**-**range PCR

Total RNAs were extracted from the human prostate tumors and the normal tissues and various cell lines using Trizol (Life Technologies). Reverse transcription was performed using the M-MLV reverse system (Takara) to obtain cDNA. qPCR was performed using with the SYBR Premix Ex Taq mix (TaKaRa) according to the manufacturer’s instructions, and the samples were run on an iQ5 Multicolor Real-time PCR Detection System (Bio-Rad). Results were normalized to expression levels for reference genes (*GAPDH* or *EIF4H*). The relative expression levels of genes were calculated using the ΔΔCT method. 3’RACE kit (Takara) was used to convert RNAs of the prostate tumors into cDNAs by a reverse transcriptase and oligo-dT adapter primer. The cDNAs were amplified by using *FOXP2*-specific primers, which annealed to exon 6, exon 11, and exon 16, and an oligo-dT adapter primer, respectively. Amplified fragments of the putative sizes were subjected to Sanger sequencing. Genomic DNAs of the fusion-positive tumors (PC_1 and PC_11) were used for long-range PCR. Amplified fragments of the putative sizes were subjected to Sanger sequencing. The primers for RT-PCR, qPCR, 3’RACE-PCR and long-range PCR were available in Table S7.

### Plasmid Construction

Constructs of *FOXP2* (Origene, #RC215021, NM_014491) and *CPED1* (Origene, #RC217158, NM_024913) were purchased. The CDS of *FOXP2*-*CPED1* fusion, truncated *FOXP2* were amplified from the corresponding fusion positive tumor and then cloned into pFlag-CMV4 vector (Sigma, #E7158). To construct the *FOXP2*-*CPED1* fusion containing the 3’UTR of *FOXP2*, the 3’UTR of *FOXP2* was generated from the *FOXP2* 3’UTR plasmid (Origene, #SC212500, NM_014491) by PCR. The PCR product was fused into *FOXP2*-*CPED1* cDNA by overlapping PCR and then the fragment of *FOXP2*-*CPED1* + 3’UTR was cloned into pFlag-CMV4 vector. To construct the plasmids for luciferase assay, the 3’UTR of *FOXP2* was amplified from *FOXP2* 3’UTR construct and ligated into a pmirGlo Dual-luciferase miRNA Target Expression Vector (Promega) to form 3’UTR-luciferase reporter vector. The full predicted seed sequences of miR-27a and miR-27b on 3’UTR of *FOXP2* were deleted from the 3’UTR-luciferase reporter vector using over-lapping PCR method to create a mutant *FOXP2* 3’UTR construct. The CDS of *FOXP2*-*CPED1* or *FOXP2* was cloned into the pCDH-CMV-MCS-EF1 Lentivector (SBI, #CD513B-1). To knockdown of endogenous *FOXP2* expression in PC3 and LNCaP cells, four shRNA fragments (1#ggaagacaatggcattaaacattcaagagatgtttaatgccattgtcttcctttttt;2# ggacagtcttcagttctaagtttcaagagaacttagaactgaagactgtcctttttt;3# gcaggtggtgcaacagttagattcaagagatctaactgttgcaccacctgctttttt;4# gcgaacgtcttcaagcaatgattcaagagatcattgcttgaagacgttcgctttttt) targeting the exon 7, exon 9 and exon 10 of *FOXP2* coding sequencing were cloned into pLent-4in1shRNA-GFP vector. The Scrambled-pLent-4in1shRNA-GFP control vector included a fully scrambled sequence. Primers used for various plasmid constructions were listed in Table S7.

### Lentivirus Transduction and Establishment of Stable Cell Lines

Lentiviruses were produced by cotransfecting the pCDH-*FOXP2* construct or pCDH-*FOXP2*-*CPED1* construct or pCDH vector or *FOXP2*-pLent-4in1shRNA-GFP vector or Scrambled-pLent-shRNA-GFP vector with the packaging plasmid Mix pPACK (SBI) in HEK293T cells. At 36 hours post-transfection, viral supernatants were collected, centrifuged at 12,000 × *g* for 20 min, filtered through 0.45 μm Steriflip filter unit (Millipore) and concentrated using ViraTrap lentivirus purification kit (Biomiga). NIH3T3 or RWPE-1 cells or PC3 or LNCaP at 90% confluence were infected with Lenti-*FOXP2*, Lenti-*FOXP2*-*CPED1*, Lenti-pCDH, Lenti-*FOXP2*-pLent-4in1shRNA-GFP and Lenti-scrambled-pLent-shRNA-GFP viruses, respectively. Cells were spun at 1,800 × *g* for up to 45 min at room temperature. After 24 hour-incubation at 37 °C, cells were split and placed into selective medium containing 1 μg/ml puromycin. Puromycin-resistant clones were grown from single cell. Western blot using anti-FOXP2 antibody (Millipore, #MABE415, 1:1,000 dilution) was performed to detect FOXP2 protein expression in *FOXP2* or *FOXP*-*CPED1*-overexpressing NIH3T3 or RWPE-1 cell clones or sh*FOXP2*-expressing PC3 or LNCaP cell clones.

### Focus Formation Assay

NIH3T3 cells stably expressing the *FOXP2*-*CPED1*, *FOXP2* and lentivirus pCDH vector, respectively, and parental cells were seeded at a concentration of 1 × 10^4^ cells (parental NIH3T3 cells were mixed with NIH3T3 cells overexpressing *FOXP2*-*CPED1*, *FOXP2* or lentiviral vector, respectively, at a 100:1 cell ratio) per well in a 6-well plate. Cells were then cultured for 14-16 days. The representative foci were either taken pictures using a Nikon microscope or fixed with 4% paraformaldehyde solution for 10 min and subsequently stained with 0.05% crystal violet and solubilized with 4% acetic acid. RWPE-1 cells stably expressing the *FOXP2*-*CPED1*, *FOXP2* and lentiviral vector, respectively, and parental cells were seeded at a concentration of 1 × 10^4^ cells per well in a 6-well plate. Cells were then cultured for 14-16 days. Then fixed with 4% paraformaldehyde solution for 10 min and subsequently stained with 0.05% crystal violet. The representative foci were taken pictures using a Nikon microscope. PC3 cells stably expressing shRNA targeting *FOXP2* or parental cells were plated at a density of 1,000 cells per well in triplicate in a 6-well plate for 10 days. LNCaP stably expressing shRNA targeting *FOXP2* cells or parental cells were plated at a density of 1,000 cells per well in triplicate in a 6-well plate for 25 days. Then fixed with 4% paraformaldehyde solution for 10 min and subsequently stained with 0.05% crystal violet.

### Soft Agar Colony Formation Assay

For the assay of NIH3T3 cells stably expressing the *FOXP2*-*CPED1*, *FOXP2*, lentiviral vector or parental cells, cells were collected and suspended in 0.4% soft agar at 1,000 cells (cell suspension in complete growth medium mixed with 2 × DMEM, 20% FBS and 1.2% agar (Difco Noble Agar, BD Bioscience, #214230) at a 1:1:1 dilution). The cell suspension (0.75 ml) was added to each well of a 24-well plate and kept at 4 °C for 20 min. For RWPE-1 cells stably expressing the *FOXP2*-*CPED1*, *FOXP2*, lentiviral vector or parental RWPE-1, cells were suspended in 0.4% soft agar at 2,000 cells (cell suspension in complete growth medium mixed with Defined Keratinocyte-SFM with 10% FBS and 1.2% agar at a 2:1 dilution). The cell suspension (0.75 ml) was added to each well of a 24-well plate and kept at 4 °C for 20 min. Each cell line was plated in quadruplicate. The fresh mediums were changed every four days and incubated for four weeks. The colony counting was performed using a Nikon microscope and representative images were then acquired.

### Cell Proliferation Assay, IC50 Assay and Drug Response Assay

For growth curve of NIH3T3 and RWPE-1 cell lines, indicated cells were seeded at 3,000 cells per well in a 96-well plate in quadruplicate, treating with increasing drug concentrations for 48 hours. Foretinib (#A2974) was purchased from ApexBio. MK2206 (#HY-10358) was purchased from MCE. The *in vitro* half-maximal inhibitory concentration (IC_50_) values were determined using Orange 8.0 software and dose response curves were plotted with package drc in R environment. For proliferation assay of RWPE-1, PC3 or LNCaP cells, indicated cells were seeded at 3,000 cells per well in a 96-well plate in quadruplicate for 6 days (RWPE-1), 8 days (PC3) and 9 days (LNCaP), respectively. The viability of cells was determined using a Celltiter-Glo assay (Promega) on INFINITE 200 Pro multimode reader (TECAN). For drug response assay, NIH3T3 cells stably expressing the *FOXP2* and *FOXP2*-*CPED1*, respectively were suspended in 0.4% soft agar at 3,000 cells (cell suspension in complete growth medium mixed with 2 × DMEM, 20% FBS and 1.2% agar at a 1:1:1 dilution) with indicated drug concentrations. Each cell line was plated in quadruplicate. The medium with indicated drug concentration was changed every four days and incubated for four weeks. The colony counting was performed using a Nikon microscope and representative images were then acquired. RWPE-1 cells stably expressing the *FOXP2* and *FOXP2*-*CPED1*, respectively were plated at a density of 3,000 cells per well in triplicate in a 6-well plate for 12 days with indicated drug concentrations. Then fixed with 4% paraformaldehyde solution for 10 min and subsequently stained with 0.05% crystal violet.

### Animal Xenograft Studies

Fifty 6 to 8-week-old non-obese diabetic severe combined immune deficiency spontaneous female mice (NOD.CB17-*Prkdc*^scid^/NcrCrl) were used. 2 × 10^6^ NIH3T3 cells in PBS were injected subcutaneously into the flanks of NOD-SCID mice (≥ 5 mice for each cell line) at an inoculation volume of 100 μl with a 23-gauge needle. Mice were monitored for tumor growth, and two months was selected as an endpoint. Tumor volumes (*V*) were calculated with the following formula: ((width)^2^ × (length))/2 = *V* (cm^3^).

### Protein Blot Analysis

Various cell lines, the frozen human tumors and normal tissues and the tumors from animal xenograft studies were lysed in RIPA lysis buffer containing protease and phosphatase inhibitors (Roche Inc), and sonicated for 20s. These extractions were resolved by SDS-PAGE and electrotransferred to PVDF membranes (Millipore). Membranes were blocked for 1 h and blotted for various primary antibodies overnight in 5% non-fat milk or 5% BSA in Tris Buffered Saline Solution, 0.5% Tween-20 (TBST) (Thermol Scientific). The following primary antibodies were used: anti-FOXP2 (Millipore, #MABE415, 1:1,000, an antibody to N terminus of FOXP2), anti-FLAG tag (M2) (Sigma, #F1804, 1:1,000), anti-Phospho-Akt (Ser473) (Cell Signalling Technology, #9271, 1:1,000), anti-Akt (Cell Signalling Technology, #9272, 1:1,000), anti-Phospho-Met (Tyr1234/1235) (Cell Signalling Technology, #3077 1:1,000), anti-Met (Abcam, #ab51067, 1:1,000), anti-Androgen receptor (Abcam, #ab133273, 1:1,000), anti-beta-Actin (C4) (Santa Cruz Technology, #sc-47778, 1:1,000), anti-alpha Tubulin (DM1A) (Abcam, #ab7291, 1:5,000). Horseradish-preoxidase conjugated antibodies to mouse (Abcam, #ab6789, 1:5,000) or to rabbit (Abcam, #ab6721, 1:5,000) were used as the secondary antibodies, and Immobilon Western Chemiluminescence HRP Substrate (Millipore) was used for detection.

### Transfection of MicroRNAs and Luciferase Reporter Assay

The microRNA mimics and microRNA inhibitors synthesized by Ribobio were used as follows: hsa-miR-19b-5p, hsa-miR-23a-3p, hsa-miR-23b-3p, hsa-miR-27a-3p, has-miR-27b-3p, hsa-miR-132-3p, hsa-miR-134-5p, hsa-miR-186-5p, hsa-miR-196b-5p, hsa-miR-212-5p, hsa-miR-214-3p, hsa-miR-379-5p, microRNA negative control (NC), hsa-miR-27a-3p inhibitor, hsa-miR-27b-3p inhibitor and negative control inhibitors (NC-I). The HEK293T cells were seeded in 96-well plate in quadruplicate and co-transfected with each of 12 microRNAs and the wild-type *FOXP2* 3’UTR firefly luciferase reporter construct using Lipofectamine 2000 (Life Technologies) for 26 hours. Wild-type or mutant *FOXP2* 3’UTR firefly luciferase reporter construct was co-transfected into HEK293T cells with NC or miR-27a or miR-27b. After 26 hours, cell lysates were prepared to consecutively measure the firefly and *Renilla* luciferase activities using Dual-luciferase reporter assay system (Promega) on INFINITE 200 Pro multimode reader (TECAN). The firefly luciferase activity was normalized by *Renilla* luciferase activity.

### Histological Analysis

Mice xenograft tumors originated from NIH3T3 cell lines which expressed *FOXP2* or *FOXP2*-*CPED1* were fixed in 4% paraformaldehyde. Consecutive paraffin sections of the mice xenograft tumors and prostates of ARR2PB-*FOXP2* or ARR2PB-*FOXP2-CPED1* transgenic mice (4 μm thickness) were used for H&E staining and immunohistochemical analyses. Specimens from 25 cases of benign prostatic hyperplasia and 45 cases of primary prostate cancer were analysed for FOXP2 staining with an immunohistochemical assay. Protein expression was evaluated as negative, low, medium and strong. The sections were pretreated with citrate buffer (pH 9.0) in a microwave oven for 20 min for antigen retrieval. The primary antibody to FOXP2 (Sigma, #HPA000382) was used at a 1:200 dilution. The following primary antibodies were used for immunohistochemical staining: anti-Ki67 (Abcam, #ab16667, 1:300), anti-Phospho-Akt (Ser473) (Cell Signalling Technology, #9271, 1:200), anti-Phospho-Met (Tyr1234/1235) (Cell Signalling Technology, #3077, 1:200).

### Pathway Enrichment Analysis

Gene set enrichment analysis of the 3206 *FOXP2* expression-correlated genes in human prostate tumors was performed by GSEA (v2.0) [26] using gene sets from Molecular Signatures Database (MSigDB) (v6.2) (http://software.broadinstitute.org/gsea/msigdb/index.jsp). Cancer-associated pathway analysis of the 3206 *FOXP2* expression-correlated genes was performed using gene sets from MSigDB and KEGG pathway database, and results were visualized using R (v3.3.1).

### Establishment of ARR2PB-*FOXP2* and ARR2PB-*FOXP2-CPED1* Transgenic Mice

To generate a mouse model expressing *FOXP2* or *FOXP2-CPED1* in a prostate-specific manner, we amplified the human *FOXP2* or *FOXP2-CPED1* cDNA sequence from the PC_1 tumor harboring the *FOXP2-CPED1* fusion by RT-PCR. A composite ARR2PB promoter [27] and a hGH_PA_terminator were introduced into the two sequence to produce ARR2PB-*FOXP2-*hGH_PA and ARR2PB-*FOXP2-CPED1*-hGH_PA by overlapping PCR method. Subsequently, they were cloned into a pBluescript SK+(PBS) expression vector that contains a T7 promoter to obtain PBS-ARR2PB-*FOXP2* and PBS-ARR2PB-*FOXP2-CPED1* using KpnI and XhoI multicloning sites. The ARR2PB*-FOXP2* and ARR2PB-*FOXP2-CPED1* mRNA were generated *in vitro* using the MEGA shortscript T7 kit and were delivered into mouse zygotes by microinjection, respectively to establish transgenic mice (C57/BL6 background). Next, mice were analyzed for construct integration by PCR genotyping assay (Table S7). In total, we identified five founder positive lines for the *FOXP2* transgene (three positive) and the *FOXP2-CPED1* transgene (two positive). Subsequently, we analysis of future offspring using PCR genotyping assay. We evaluated expression of human *FOXP2* and *FOXP2-CPED1* at the mRNA and protein levels by RT-PCR, western blot and immunohistochemical staining. All the five founder lines showed mRNA and protein expression of human *FOXP2* and *FOXP2-CPED1* in all lobes of the transgenic mice prostate. Founder lines 16^#^ and 48^#^ for the *FOXP2* transgene and founding line 26^#^ for the *FOXP2-CPED1* transgene were subsequently used for phenotypic analysis.

### Statistics

We used SPSS (v16.0), Origin (v8.0) or R (v3.3.1) (The R Project for Statistical Computing, http://www.r-project.org/) software for statistical calculation. Specific statistical tests, number of samples, experimental or public data utilized in each analysis were shown along the main text or in the figure legends. Cell culture-based experiments were conducted three times or more, which were triplicated or quadruplicated. The data are presented as mean ±s.d. Student’s *t* test was used to compare the difference between means in normal distributions. In the boxplots, boxes display the 25th to 75th percentiles, lines represent the medians, and whiskers represent 1.5 × the interquartile range. Mann-Whitney *U* test was used to compare the difference between data in non-normal distributions. The two-tailed *P* < 0.05 was considered to be statistically significant. Spearman rank correlation was used to measure the association between expressions of individual genes. We used the Cochran-Mantel-Haenszel test to estimate adjusted odds ratio (OR_MH_) and adjusted *P* values when *ETS*-fusions were considered a confounding factor.

## Results

### Enhanced *FOXP2* Expression in Prostate Cancer Plays an Oncogenic Role

*FOXP2* mutations have long been thought to be the cause of developmental speech and language disorder in humans [9–11]. In this study, we identified a new gene fusion, *FOXP2-CPED1*, in 2 of 100 indolent prostate tumors by performing RNA sequencing and whole-genome sequencing analyses (Fig. S1A-S1F and Table S1-S2) (see details in the Supplementary Results). We found that in the prostate tumor, the fusion consequently resulted in increased expression of a truncated FOXP2 protein that retained the complete FOXP2 functional domains [11, 28–30], but had an aberrant C-terminus (Fig. S1H-S1K) (see details in the Supplementary Results). Mechanistically, we conducted small RNA sequencing of the fusion-positive tumor followed by functional assays, demonstrating that *FOXP2-CPED1* fusion led to loss of the 3’UTR of *FOXP2*, thus allowing escape from regulation by miR-27a and miR-27b and consequently resulting in aberrantly high expression of the truncated FOXP2 protein with full functional domains (Fig. S1L-S1P) (see details in the Supplementary Results).

To date, the *FOXP2* expression pattern in prostate cancer has remained unclear. We therefore evaluated the expression of *FOXP2* in prostate cancer by analysing several datasets, including our in-house samples (primary prostate cancer tumors, n = 92; matched normal tissues, n = 23), the SU2C dataset (metastatic prostate cancer samples, n = 117) [31] and the GTEx dataset (normal prostate tissues, n = 106). The *FOXP2* mRNA levels were significantly increased in prostate cancer samples with respect to normal tissues (Fig. 1A-1B). We then examined tissue extracts by immunoblotting. The amounts of FOXP2 protein were evidently increased in primary prostate adenocarcinomas relative to the matched normal tissues and benign prostatic hyperplasia (BPH) samples (Fig. S2A-S2B). We further carried out an immunohistochemical assay on 25 BPH samples and 45 primary prostate tumors. Strong or medium staining of FOXP2 protein was observed in 58% (26/45) of primary prostate adenocarcinomas. No or low nuclear staining of FOXP2 was observed in 96% (24/25) of benign prostatic tissues. There was a statistically significant difference in the FOXP2 staining intensity between prostate cancer and benign prostate tissue (two-tailed *P* = 0.001, by Fisher’ exact test) (Fig. 1C). Likewise, *FOXP2* expression was markedly elevated in human prostate cancer cell lines (PC3, LNCaP, VCaP and DU145), but was undetectable or low in normal or immortalized human prostate epithelial cells (HPrEC and RWPE-1) (Fig. 1D-1E). Given the data above, these findings suggested that there is a tendency for increased expression of FOXP2 protein between normal tissue and prostate neoplasia. Together, our data indicated that *FOXP2* was highly expressed in human prostate tumors.

**Figure 1.**
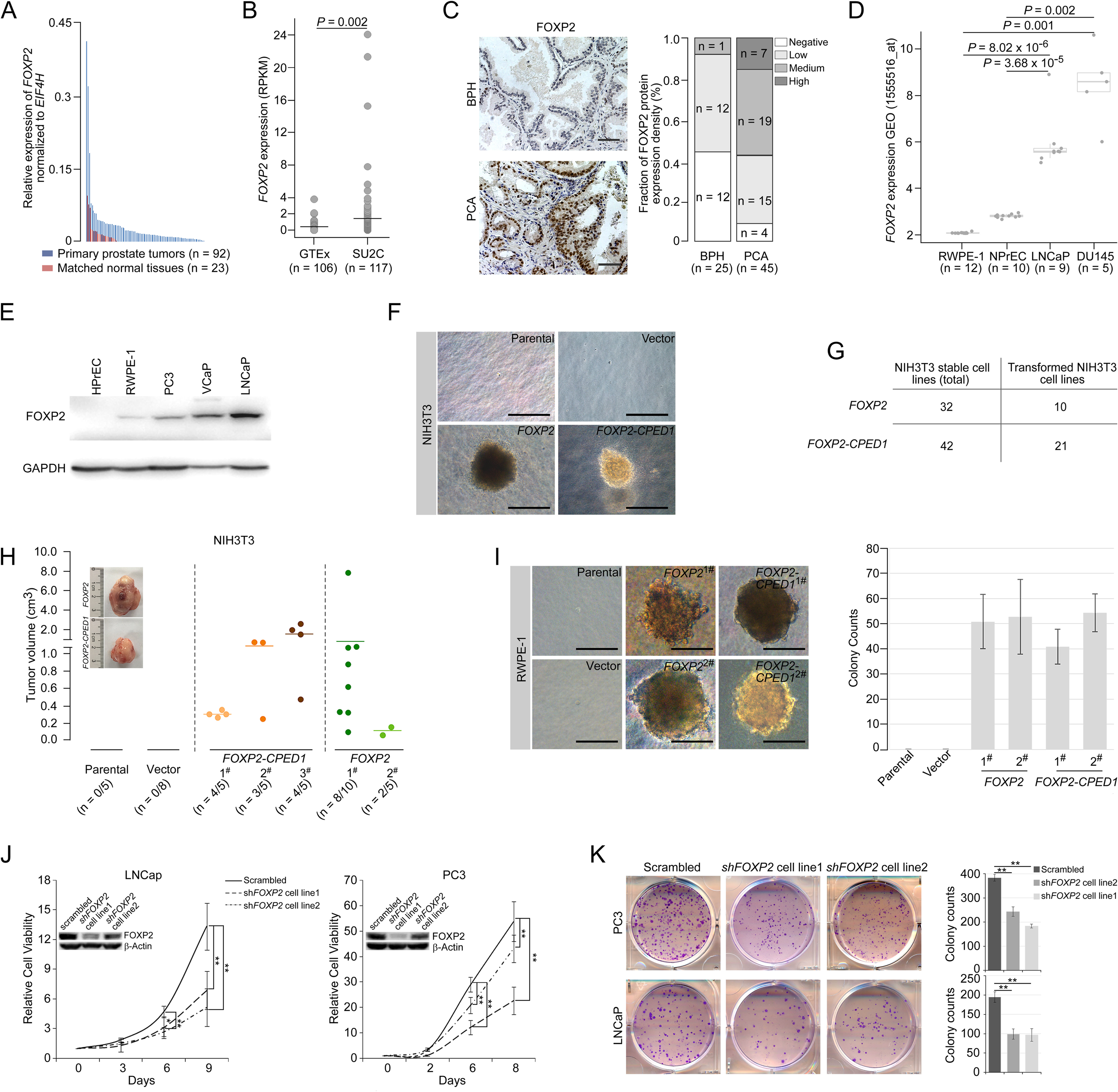
*FOXP2* functions as an oncogene in prostate cancer. **A.** Expression of *FOXP2* mRNA in 92 primary prostatic adenocarcinoma tissues and 23 matched normal tissues analysed by qPCR. **B.** Expression of *FOXP2* mRNA in normal prostate samples from the GTEx dataset (n = 106) and metastatic prostate tumors from the SU2C dataset (n = 117). The horizontal bar indicates the mean in each group. *P* values were calculated by 2-tailed Mann-Whitney *U* test. **C.** *Left*, Representative images of FOXP2 protein expression in benign prostatic hyperplasia (BPH) (n = 25) or primary prostatic adenocarcinoma (PCA) (n = 45) by immunohistochemistry. Scale bars, 100 μm. *Right*, The bar chart indicates the number of PCA and BPH with negative, low, medium and high FOXP2 expression, respectively. **D.** Expression of *FOXP2* mRNA in two types of human benign prostate epithelial cells, RWPE-1 and NPrEC, and in two prostate cancer cell lines LNCaP and DU145 from the GEO database (1555516_at). *P* values were calculated by 2-tailed Student’s *t* test. **E.** Western blot measuring FOXP2 protein in one normal human prostate epithelial cell HPrEC and one benign prostate epithelial cell RWPE-1, and in three human prostate cancer cell lines, PC3, VCaP and LNCaP. The experiment was repeated twice with similar results. **F.** Cell transformation and anchorage-independent growth were measured by a soft agar assay in parental NIH3T3 cells and NIH3T3 cells stably expressing the empty vector, *FOXP2* and *FOXP2*-*CPED1*. Scale bars, 100 μm. See Fig. S2F for details. **G.** Table indicating the number of total NIH3T3 cell lines overexpressing *FOXP2* (n = 32) or *FOXP2*-*CPED1* (n = 42) tested by soft agar colony formation assay (left column) and the number of transformed NIH3T3 cell lines induced by *FOXP2* (n = 10) or *FOXP2*-*CPED1* (n = 21) (right column). See Fig. S2F for details. **H.** Quantification of the tumor volume in NOD-SCID mice injected with NIH3T3 cells stably expressing vector, *FOXP2* (two cell lines) or *FOXP2*-*CPED1* (three cell lines) for 8 weeks. *Inset*, Representative images of xenografted tumors derived from the corresponding cells. See Fig. S2G-S2H for details. **I.** *Left*, Soft agar assay showing that RWPE-1 cells stably expressing *FOXP2* (two cell lines) or *FOXP2*-*CPED1* (two cell lines) were able to form colonies. Scale bars, 100 μm; *Right*, Quantitative analysis of the colony formation ability of the corresponding RWPE-1 cells. Mean ± s.d.; n = 4. **J.** Cell growth of prostate cancer cell lines (LNCaP and PC3) stably expressing control vector or *FOXP2* shRNA (two clones, #1 and #2). \**P* < 0.05, \*\**P* < 0.005 by 2-tailed Student’s *t* test, mean ± s.d.; n = 4. *Inset*, Immunoblot showing knockdown of FOXP2 protein in LNCaP or PC3 cells. **K.** *Left*, Focus formation assay performed with PC3 or LNCaP cells stably expressing control vector or *FOXP2* shRNA (two clones, #1 and #2). *Right*, Bar graph showing the number of colonies formed by PC3 or LNCaP cells after *FOXP2* silencing. \*\**P* < 0.005 by 2-tailed Student’s *t* test; mean ± s.d.; n = 4.

Overexpression of specific genes such as *ERG* [6, 32] and *EZH2* [33] results in a phenotype of hyperplasia and promotes of invasive properties in the prostate, indicating a critical proto-oncogenic role in prostate tumorigenesis. Therefore, in this study, to explore the biological consequence of high expression of the *FOXP2* gene, we first introduced the wild-type or fusion *FOXP2* cDNA into mouse NIH3T3 fibroblasts that lack endogenous Foxp2 protein expression (Fig. S2C-S2D) and then carried out a colony formation assay. NIH3T3 fibroblasts, as normal fibroblasts, are considered to be entirely anchorage-dependent for proliferation. We found that *FOXP2* can transform NIH3T3 fibroblasts, inducing loss of contact inhibition and gain of anchorage-independent growth, which are tumorigenic properties (Fig. 1F-1G and Fig. S2E-S2F). We also noted that, as anticipated from the structural properties of the *FOXP2*-*CPED1* fusion, it was also able to confer this oncogenic phenotype as well (Fig. 1F-1G and Fig. S2E-S2F). We next determined the oncogenic activity of *FOXP2* and its rearrangement lesion in a NOD-SCID mouse model. We observed that *FOXP2*-overexpressing NIH3T3 cells injected into the flanks of NOD-SCID mice formed tumors at a higher penetrance than did parental or empty vector control cells: in 10/15 mice compared with 0/5 mice for parental cells and 0/8 for control cells (Fig. 1H and Fig. S2G-S2H). In 11/15 mice, *FOXP2*-*CPED1*-overexpressing NIH3T3 cells formed tumors in mice (Fig. 1H and Fig. S2G-S2H).

The oncogenic properties of *FOXP2* were further shown by its ability to transform a human prostate epithelial cell line, RWPE-1, which is immortalized but nontransformed [34]. We observed that *FOXP2* overexpression significantly increased the proliferation of RWPE-1 cells relative to that of control cells (Fig. S2I). Similar to NIH3T3 cells, assays of focus formation and anchorage-independent growth consistently showed that overexpression of wild-type *FOXP2* or *FOXP2* fusion induced malignant transformation of RWPE-1 cells (Fig. 1I and Fig. S2J). Conversely, we carried out *FOXP2* short hairpin RNA (shRNA) knockdown in two prostate cancer cell lines (LNCaP and PC3). We found that shRNA-mediated *FOXP2* knockdown markedly inhibited the cell growth and colony-forming ability of LNCaP and PC3 cells (Fig. 1J-1K). Our findings were consistent with previous data showing that the cell growth ability was decreased by *Foxp2* knockdown in mouse pancreatic cancer cells [35]. Taken together, our data demonstrated that *FOXP2* is tumorigenic in prostate cancer.

### *FOXP2* Overexpression Aberrantly Activates Oncogenic MET Signalling

To explore the molecular mechanism underlying the role of *FOXP2* in prostate cancer, we first carried out analysis of the entire expression spectrum of 255 primary prostate tumors from TCGA (Fig. 2A). We observed that the expression of *FOXP2* was significantly correlated with the expression of 3206 genes (refer to *FOXP2* expression-correlated genes, FECGs) (|Spearman’s rho ≥ 0.5|) (Fig. 2B and Table S3). Gene set enrichment analysis (GSEA) of these FECGs identified that the top significantly enriched gene sets corresponded to 11 known prostate cancer gene sets, including Wallace_Prostate_Cancer_Race_Up, Acevedo_FGFR1_Targets_In_Prostate_Cancer_Model, Kondo_Prostate_Cancer_with_H3k27me3 and Kras.Prostate_Up.V1_DN signatures (Fig. 2C and Table S4). Pathway enrichment analysis identified that Dairkee_Tert_Targets_Up, PI3K-AKT Signalling, Mtiotic_Spindle, TGF_Beta_Signalling and Inflammatory_Response were significantly enriched in positively *FOXP2*-correlated genes. E2F_Targets, DNA_Repair and Oxidative_Phosphorylation were significantly enriched in negatively correlated genes (Fig. 2D and Table S5). Notably, 18 of the 74 FECGs enriched in the PI3K-AKT pathway are known cancer driver genes, which include 4 core members of oncogenic MET signalling (*HGF*, *MET*, *PIK3R1* and *PIK3CA*) (Fig. 2D).

**Figure 2.**
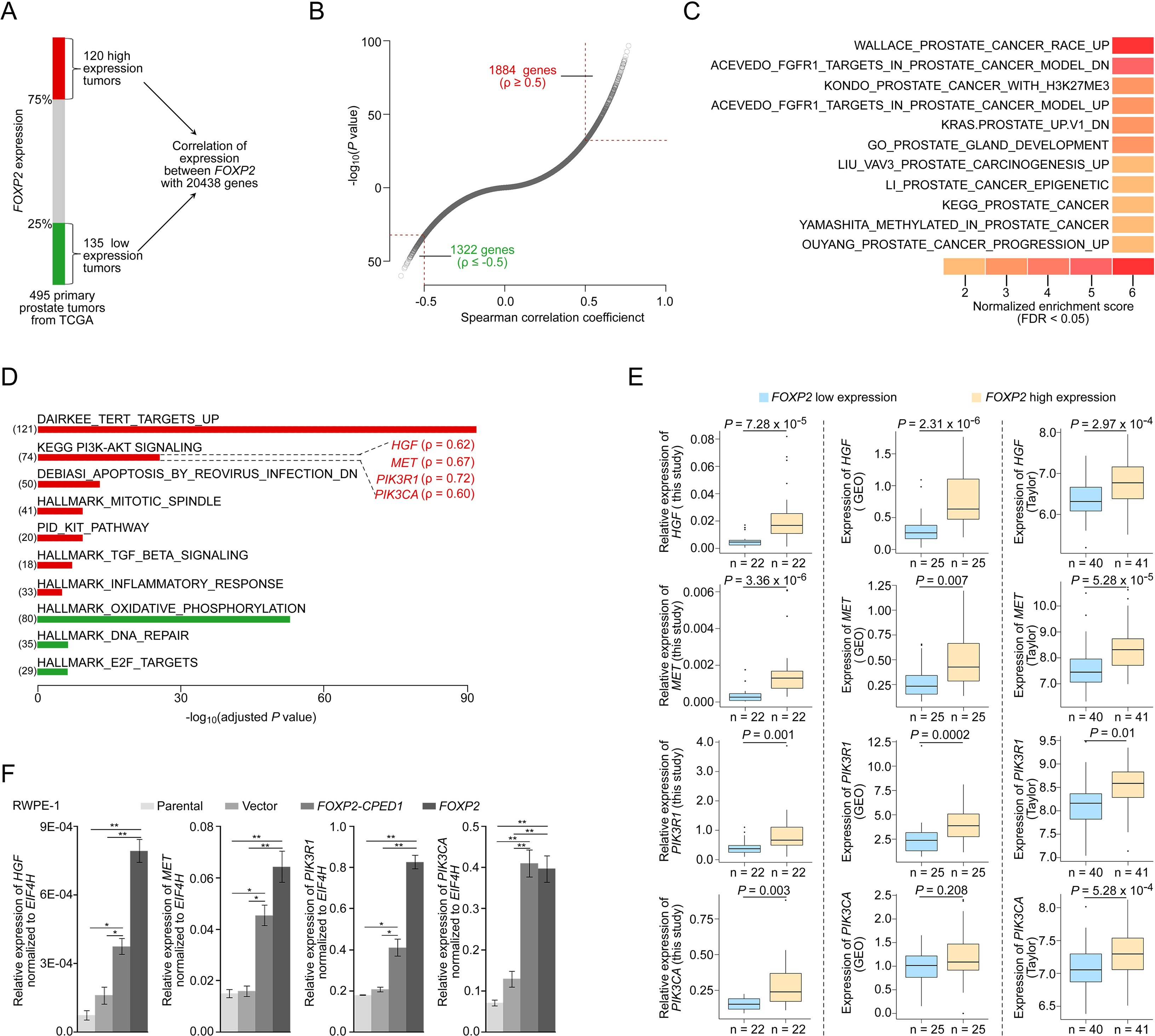
*FOXP2* activates oncogenic MET signaling in *FOXP2-*overexpressing cells and prostate tumors. **A.** Schematic indicating the analysis of the TCGA dataset (Prostate Adenocarcinoma, Provisional, n = 495). In total, n = 255, top 25%, high expression (n = 120), bottom 25%, low expression (n = 135). **B.** In the 255 primary prostate tumors shown in (**A**), the expression of 3206 genes was significantly correlated with the expression of *FOXP2* (|Spearman’s ρ ≥ 0.5|; 1884 significantly positively correlated genes; 1322 significantly negatively correlated genes). See also Table S3. **C.** The significant enrichment of the 3206 *FOXP2* expression-correlated genes (FECGs) in known prostate cancer gene sets from the Molecular Signatures database by GSEA. *P* values were calculated by 2-tailed Fisher’s exact test. See also Table S4. **D.** Analysis of the biological pathways of 3206 *FOXP2* expression-correlated genes using gene sets from MSigDB and KEGG. The number in parentheses corresponds to *FOXP2* expression-correlated genes that are enriched in the corresponding pathway. The Spearman correlation coefficients between the expression of *FOXP2* and the expression of four core components of MET signaling (*HGF*, *MET*, *PIK3R1* and *PIK3CA*) are indicated in parentheses. Two-tailed *P* values by Fisher’s exact test and adjusted by Bonferroni correlation. See also Table S5. **E.** The correlation analysis of gene expression between *FOXP2* and MET signalling members in our primary prostate tumors (n = 92) and two other human primary prostate cancer datasets (GSE54460, n = 100; Taylor, n = 162) by qPCR. *HGF*, *MET*, *PIK3R1* and *PIK3CA* expression levels (normalized to *EIF4H*) classified by *FOXP2* expression level (bottom 25%, low expression; top 25%, high expression). *P* values were calculated by 2-tailed Mann-Whitney *U* test. **F.** In *FOXP2*- or *FOXP2*-*CPED1*-transformed RWPE-1 cells and control cells (parental cells and empty vector-expressing cells), the relative mRNA expression levels of *HGF*, *MET*, *PIK3R1* and *PIK3CA* (normalized to *EIF4H*) were examined by qPCR. *P* values calculated by 2-tailed Student’s *t* test, mean ± s.d.; n = 3. \**P* < 0.05, \*\**P* < 0.005.

We further examined the correlation between the MET pathway and *FOXP2* in three additional datasets including our primary prostate cancer data, GSE54460 [36] and Taylor [37]. Consistently, we observed a significantly positive correlation between the expression of *FOXP2* and the four individual core members of MET signalling (*HGF*, *MET*, *PIK3R1* and *PIK3CA*) across the three datasets (Fig. 2E). Previous studies have reported that the receptor tyrosine kinase MET and its ligand HGF are important for the growth and survival of several tumor types, including prostate cancer [38, 39]. These four putative candidates were further confirmed to be upregulated in RWPE-1 cells overexpressing *FOXP2* by qPCR (Fig. 2F). A similar effect was also observed in NIH3T3 cell lines that ectopically expressed *FOXP2* (Fig. S3). These data strongly suggested that oncogenic MET signalling is activated by *FOXP2* in prostate tumors.

### Targeting MET signalling inhibits *FOXP2-*induced oncogenic effects

We next assessed whether activation of MET signalling plays a crucial role in *FOXP2*-driven oncogenic effects. We observed obviously elevated phosphorylation of tyrosines Y1234/1235 in the MET kinase domain in the *FOXP2*-transformed RWPE-1 and *FOXP2*-transformed NIH3T3 cells (Fig. 3A). Furthermore, overexpression of *FOXP2* induced strong activation of PI3K signalling, as indicated by the elevated phosphorylation level of AKT at serine 473 (Fig. 3A). AKT is a key downstream effector of HGF/MET/PI3K signalling [40] and is able to initiate prostate neoplasia in mice [41]. Conversely, FOXP2 silencing using shRNA caused decreases in phospho-MET and phospho-AKT levels in two prostate cancer cell lines, PC3 and LNCaP (Fig. 3B). Moreover, we detected higher phosphorylation levels of MET and AKT in primary human prostate tumors with higher compared with lower FOXP2 protein abundance (Fig. 3C). Since the *FOXP2*-overexpressing cells exhibited an increase in p-MET/p-AKT levels, we hypothesized that MET and AKT inhibitors could have a therapeutic benefit. The *FOXP2*-overexpressing cells were subsequently treated with the MET tyrosine kinase inhibitor foretinib or AKT inhibitor MK2206, resulting in twofold-threefold (for foretinib) and threefold-twelvefold (for MK2206) decreases in the half-maximal inhibitory concentration (IC_50_) in RWPE-1 cells and NIH3T3 cells with *FOXP2* overexpression (Fig. 3D-3E). Furthermore, treatment of *FOXP2*-overexpressing RWPE-1 cells and *FOXP2*-overexpressing NIH3T3 cells with foretinib or MK2206 abrogated MET and/or AKT phosphorylation (Fig. 3F-3G) and resulted in significantly reduced anchorage-independent growth or foci formation relative to those in nontreated cells (Fig. 3I-3H). These results demonstrated the involvement of MET signalling in the cellular transformation driven by *FOXP2*.

**Figure 3.**
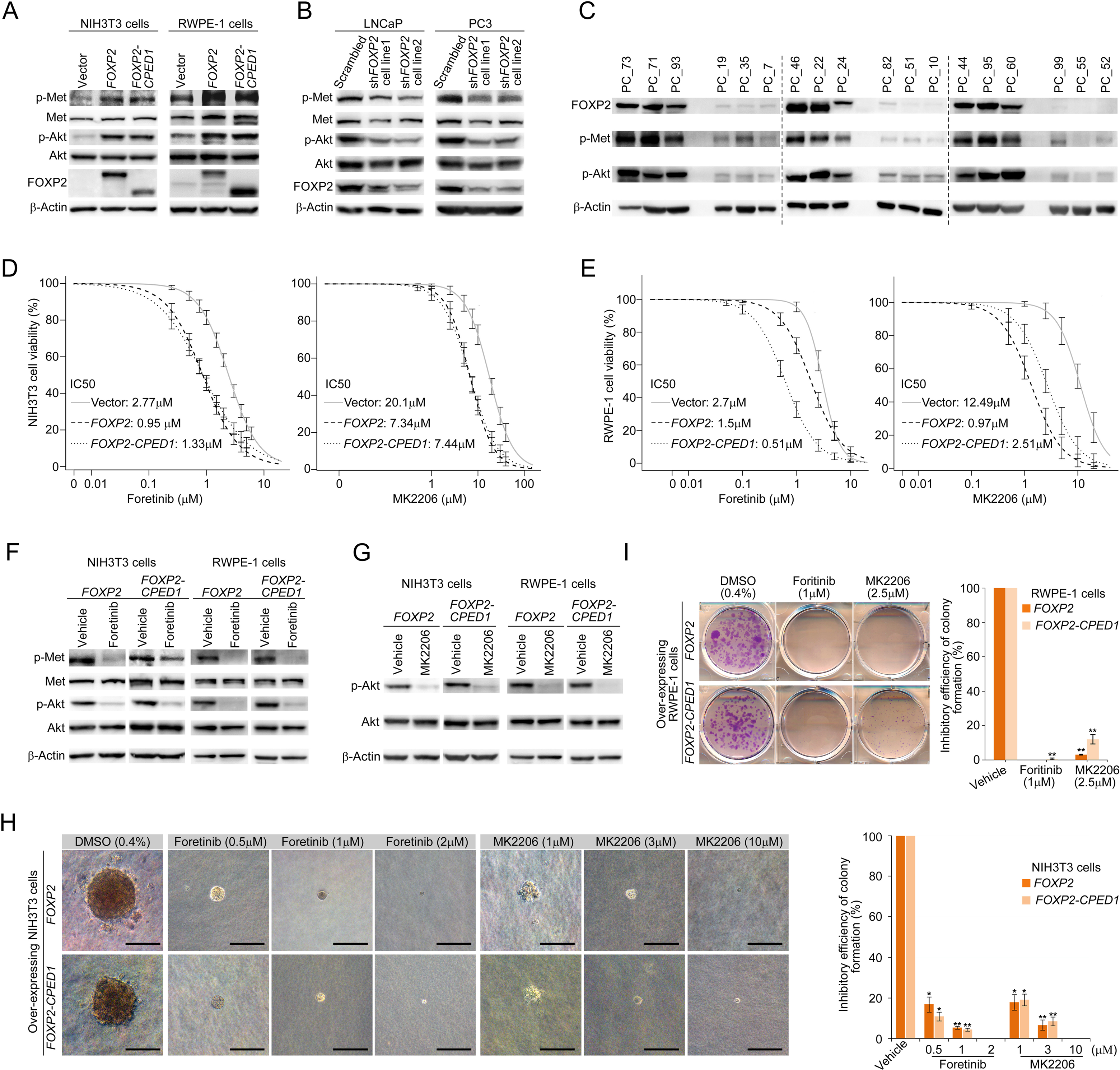
Targeting MET signaling inhibits *FOXP2-*induced oncogenic effects. **A.** Immunoblotting of the expression levels of p-Met (Y1234/1235) and p-Akt (S473) in NIH3T3 cells overexpressing *FOXP2* or *FOXP2*-*CPED1* (*left*) or in RWPE-1 cells overexpressing *FOXP2* or *FOXP2*-*CPED1* (*right*). The experiment was repeated twice with similar results. **B.** Protein blot analysis of the expression of p-Met (Y1234/1235) and p-Akt (S473) in LNCaP (*left*) or PC3 cells (*right*) that stably expressed scrambled vector and *FOXP2* shRNA, respectively. The experiment was repeated twice with similar results. **C.** Protein blot analysis of the activity of MET (Y1234/1235) and AKT (S473) in human localized primary prostate tumors (n = 18) with high and low FOXP2 expression, respectively. **D-E**. IC50 curves of the vector, *FOXP2*- or *FOXP2*-*CPED1*-overexpressing NIH3T3 cells (**D**) or -overexpressing RWPE-1 cells (**E**) treated with the MET inhibitor foretinib or the AKT inhibitor MK2206 assessed at 48 h after the treatment. Mean ± s.d.; n = 3. **F-G**. Western blot analysis of the expression levels of p-Met (Y1234/1235) and p-Akt (S473) in the *FOXP2*- or *FOXP2*-*CPED1*-overexpressing NIH3T3 cells or -overexpressing RWPE-1 cells after treatment with vehicle (0.4% DMSO) and the MET inhibitor foretinib (0.5 μM for NIH3T3 cells, 1 μM for RWPE1 cells) (**F**) or the AKT inhibitor MK2206 (1 μM for NIH3T3 cells, 2.5 μM for RWPE1 cells) (**G**). **H**. *Left*, Soft agar assay of the stable *FOXP2-* or *FOXP2*-*CPED1*-overexpressing NIH3T3 cells treated with vehicle, MET inhibitor or AKT inhibitor. Images of representative cells treated with the MET inhibitor foretinib and AKT inhibitor MK2206. Scale bars, 100 μm. *Right*, Bar graph showing the percentage of the colonies formed by these NIH3T3 cells after treatment with the indicated MET or AKT inhibitors normalized to those of DMSO-treated cells. \**P* < 0.01, \*\**P* < 0.005 by 2-tailed Student’s *t* test, mean ± s.d.; n = 4. **I**. *Left*, Focus formation assay of the stable *FOXP2-* or *FOXP2*-*CPED1*-overexpressing RWPE-1 cells treated with vehicle, the MET inhibitor foretinib or the AKT inhibitor MK2206. *Right*, Bar graph showing the percentage of the colonies formed by these REPW-1 cells after treatment with the indicated MET or AKT inhibitors normalized to those of DMSO-treated cells. \*\**P* < 0.005 by 2-tailed Student’s *t* test, mean ± s.d.; n = 4.

### Prostate-specific overexpression of *FOXP2* causes prostatic intraepithelial neoplasia (PIN)

To further determine the effects of *FOXP2 in vivo*, we generated mice with prostate-specific expression of *FOXP2* and the *FOXP2-CPED1* fusion under the control of a modified probasin promoter regulated by androgen [27] (Fig. S4A-S4E). By 46-65 weeks of age, all ARR2PB-*FOXP2* mice (n = 35) and ARR2PB*-FOXP2-CPED1* mice (n = 32) examined had hyperplasia in prostates (Table S6). Furthermore, 97% (34/35) of *FOXP2* transgenic mice and 88% (28/32) of *FOXP2-CPED1* mice developed PIN. All littermate wild-type prostates had normal morphology, although they also had focal benign hyperplasia. Most PINs were observed in the anterior and ventral prostate lobes, and they were all focal (Table S6). The foci of the lesion had multiple layers of atypical epithelial cells and presented papillary, tufting or cribriform patterns. The atypical cells exhibited poorly oriented and markedly enlarged nuclei with severe pleomorphism, hyperchromasia and prominent nucleoli (Fig. 4A and Fig. S4F). In comparison with wild-type control mice, both transgenic *FOXP2* and *FOXP2-CPED1* fusion mice showed a 5- to 6-fold increase in the proliferation index of prostate epithelial cells in preneoplastic prostate glands, as shown by Ki67 staining (Fig. 4B). Moreover, consistent with our *in vitro* data, increased phosphorylation levels of mouse Met and its downstream mediator Akt in prostates of both *FOXP2* and *FOXP2-CPED1* fusion transgenic mice compared with those in prostates of control mice were observed (Fig. 4C). Together, our data indicated that *FOXP2* has an oncogenic role in prostate tumorigenesis.

**Figure 4.**
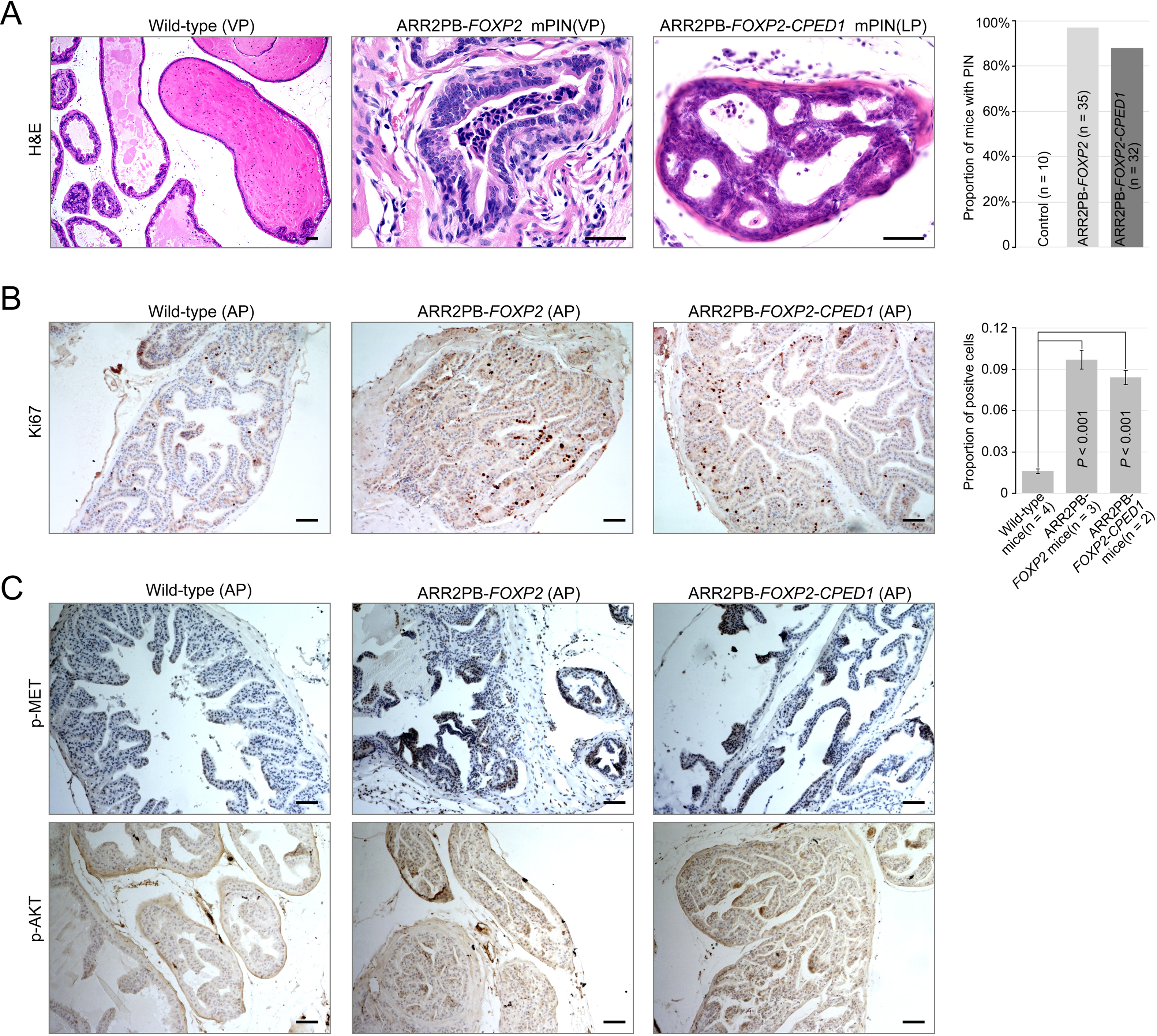
Prostate-specific overexpression of *FOXP2* causes prostatic intraepithelial neoplasia. **A.** Histological images of prostatic intraepithelial neoplasia in ARR2PB-*FOXP2* or ARR2PB-*FOXP2-CPED1* mice (200×) compared to wild-type control (40×) (mice 14 months of age). A bar graph showing the incidence of mPIN in transgenic mice at an average of 14 months of age. See Table S6 for details. **B.** Immunohistochemistry of Ki67 in prostate glands reveals a significant increase in proliferation for ARR2PB-*FOXP2* or ARR2PB-*FOXP2-CPED1* mice at 6 months of age (100×). The bar graph shows the proportion of Ki67-positive cells per gland (mean ± s.d.) reported for at least three representative prostate glands per mouse. *P* values were calculated by 2-tailed Student’s *t* test. **C.** Immunohistochemical analysis of Met (Y1234/1235) (*upper*) and Akt (S473) (*lower*) activity in prostate glands shows the upregulation of Met signaling in ARR2PB-*FOXP2* and ARR2PB-*FOXP2-CPED1* mice.

## Discussion

In this study, we determined for the first time that *FOXP2* is an oncogene in prostate tumor and that the overexpressed FOXP2 protein causes malignant transformation of normal prostate epithelial cells in humans and mice. *FOXP2* encodes an evolutionally conserved forkhead box transcription factor and is highly expressed in human thyroid, lung and smooth muscle tissue; it is particularly highly expressed in the brain. However, the *FOXP2* expression pattern in normal prostate tissue and malignant neoplasms of the prostate has not been clearly characterized. Our findings revealed that FOXP2 protein expression is absent or low in normal human prostate epithelial cells and benign prostatic hyperplasia but is markedly increased in prostatic intraepithelial neoplasia, localized prostate tumors and metastatic prostate cancer cell lines. We found that primary prostate tumors with *FOXP2* expression had a higher Gleason grade than cases without *FOXP2* expression by analysing the clinical data of 491 human TCGA primary prostate tumors (Fig. S5A). Moreover, we observed frequent amplification of the *FOXP2* gene, which occurred in 15-20% of primary prostate tumors from the Broad/Cornell [42] and TCGA datasets and in 25% of metastatic tumors from the SU2C dataset [31] (Fig. S5B). Frequent *FOXP2* gain also occurred in 18 other types of human solid tumors (Fig. S5C). Finally, we evaluated the clinical significance of *FOXP2* copy number alterations (CNAs) in 487 primary prostate tumors. CNAs of *FOXP2* were significantly associated with high Gleason scores (Gleason score ≥ 8) (OR = 2.10; 95% CI, 1.38-3.22; *P* = 0.001, by Fisher’s exact test) and high-grade pathologic T stages (T ≥ 3a) (OR = 2.01; 95% CI, 1.26-3.21; *P* = 0.003, by Fisher’s exact test) (Fig. S5D). These data suggested that the genomic lesion in *FOXP2* might contribute to high-risk prostate cancer. In addition, we found that CNAs of *FOXP2* were prominently enriched in ETS fusion-negative prostate tumors from the MSKCC/DFCI dataset (n = 685) [43] (*P* = 2.42 × 10^-6^, by Fisher’s exact test) (Fig. S5E), suggesting that *FOXP2* CNAs and ETS fusions were partially mutually exclusive. Similar to our observation, Stumm *et al*. reported moderate to strong FOXP2 protein expression in 75% of prostate tumors, and a higher protein expression level of FOXP2 was correlated with higher Gleason score, advanced T stage and earlier cancer recurrence in *ERG* fusion-negative prostate cancers [44].

Here, we identified a novel recurrent *FOXP2-CPED1* fusion in two localized prostate tumors. Due to loss of miR-27a/b-mediated transcriptional regulation, the fusion yielded a truncated FOXP2 protein that was highly expressed in the fusion-carrying tumor (Fig. S1H-S1P). In addition, whole-genome sequencing of the fusion-positive tumor suggested that the *FOXP2* fusion was an early event in the tumor (Fig. S1G). The truncated FOXP2 protein retained the full forkhead DNA binding domain (DBD). In humans, mutations in the DBD of FOXP2 cause severe autosomal dominant disorders of speech and language [29]. Several lines of evidence also showed that the DBD of some other FOX family proteins was crucial for the growth and invasion of tumors [45, 46]. Consistent with the observation of cellular transformation resulting from *FOXP2* overexpression, ectopically expressed *FOXP2-CPED1* induced the transformation of normal or benign prostate cells to malignant cells *in vitro* and *in vivo* and aberrantly activated oncogenic MET signalling, suggesting that the oncogenic role of the FOXP2 protein could be DBD dependent. Taken together, these findings for the *FOXP2* fusion further supported that *FOXP2* was an oncogene implicated in the tumorigenicity of the prostate.

To identify genes that are significantly regulated by *FOXP2*, we compared two sources of differential gene expression data from TCGA primary prostate tumors and our *FOXP2*-transformed NIH3T3 cells (Fig. S3E). We found 60 common genes whose expression was significantly correlated with *FOXP2* expression. The overlapping genes included five known cancer driver genes (including *HGF*, *MET* and *PIK3R1*) (Fig. 2D and Fig. S3C-S3D) and five putative *FOXP2* targets (Fig. S3E), such as *MET* and *PIK3R1*, identified by chromatin immunoprecipitation assays in both mice and humans [11, 15, 47–49].

Immunohistochemical staining of normal and malignant human prostate samples showed that MET protein expression is absent in epithelial cells of normal prostate glands and low in benign prostate hyperplasia, whereas it is frequently detected in PIN, localized and metastatic prostate tumors [50, 51]. Transgenic overexpression of Met in mouse prostate epithelial cells is sufficient to induce PIN under HGF conditions and markedly promotes prostate tumor initiation, invasion and metastasis in a Pten-deficient background [52]. Our data showed that overexpression of *FOXP2* activated oncogenic MET signalling in *FOXP2*-transformed prostate epithelial cells and human prostate tumors, and inhibition of MET signalling activation in *FOXP2*-overexpressing RWPE-1 cells and NIH3T3 cells significantly suppressed the anchorage-independent growth or foci formation of these cells. In agreement with our findings, treatment of both PC3 and LNCaP prostate tumor cells with a MET inhibitor resulted in reduced cell proliferation [53]. Together, these data indicated that *FOXP2* is implicated in the initiation and progression of prostate cancer by aberrantly activating MET signalling.

Phase III clinical trials have shown that the MET kinase inhibitor cabozantinib has some clinical activity in patients with advanced prostate cancer, including conferring improvements in the bone scan response, radiographic progression-free survival, and circulating tumor cell conversion, although it failed to significantly improve overall survival [54, 55]. Qiao *et al.* provided experimental evidence that MET inhibition was only effective in prostate cancer cells with MET activation [56]. Our data showed that overexpressed *FOXP2* aberrantly activated oncogenic MET signalling in transformed prostate epithelial cells and human prostate tumors. We observed that *FOXP2*-overexpressing cells were more sensitive to inhibitors of the MET signalling pathway than control cells. Collectively, these data suggested a potential therapeutic option for prostate cancer patients with high expression levels of *FOXP2*. Future work is required to determine whether additional pathways are activated or inhibited by *FOXP2*. In summary, our data demonstrated that *FOXP2* functions as an oncogene involved in tumorigenicity of prostate. Aberrant *FOXP2* expression activates MET signalling pathway in prostate cancer with potential therapeutic implication.

## Declarations

### Ethics approval and consent to participate

This study was approved by the Research Ethics Board of the Beijing Hospital, National Health Commission and performed in accordance with the relevant guidelines and regulations. All participants provided written informed consent before participation. Animal procedures were approved by the Institutional Animal Care and Use Committee of National Institute of Biological Sciences, Beijing (Beijing, China).

### Consent for publication

Not applicable.

### Data and Software Availability

The RNA-Seq data from 10 pairs of primary prostate tumors and normal tissues has been deposited in GEO database with the accessions codes: GSE114740. The Whole Genome Sequencing data from *FOXP2*-*CPED1* fusion-positive tumor (PC_1) has been deposited in SRA database with the accessions: SRR7223723. All other available public data supporting findings of this study can be found in the manuscript or its supplementary files obtained from the cBioPortal database (http://www.cbioportal.org/), the GTExPortal database (https://gtexportal.org/home/documentationPage) and the Gene Expression Omnibus database (https://www.ncbi.nlm.nih.gov/geoprofiles). We considered the prostate cancer samples with *FOXP2* mRNA expression level < 0.5 normalized RNA-seq by expectation maximization (RSEM) from the TCGA dataset (Prostate Adenocarcinoma, Provisional) as *FOXP2*-negative tumors. **URLs**. Rfam database, http://rfam.xfam.org/; NCBI Genbank, http://www.ncbi.nlm.nih.gov/genbank/; IREAP, https://sourceforge.net/projects/mireap/; miRNA reference sequences, http://www.mirbase.org/, release 19; Break Dancer, http://breakdancer.sourceforge.net/; TargetScan, http://www.targetscan.org/vert_71/; microRNA.org, http://www.microrna.org/; PICTAR5, http://pictar.mdc-berlin.de/; Cancer Gene Census, https://cancer.sanger.ac.uk/census.

## Author contributions

X. Zhu, Y. Zhao, J.W., Z.Y. developed the concept and designed research. X. Zhu, Y. Zhao, C.C., J.W., W.J., S.L., Y.L., F.W. and R.Y. performed experiments or bioinformatics analyses. X. Zhu, Y. Zhao, C.C. and L.P. analysed the data. W.Z., X.Y performed histological analyses. D.W., Y.X., J.Z., J.L., Y.Q., Y. Zhang, P.W., Q.H., L.Z. and R.Q. contributed samples. X. Zhu and Y. Zhao wrote the manuscript. All authors discussed the data and made the manuscript revised. J.W., Z.Y. X., X. Zhu and Y. Zhao jointly supervised this work. All authors read and approved the final manuscript.

## Acknowledgments

We thank Mingming Shi, Qiang Gao and Xuanmin Guang for helpful advice and assistance with analyses.

## Supplementary Results

We identified one previously uncharacterized out-of-frame fusion involving *FOXP2* and *CPED1* in 2 of 100 tumors by performing RNA-Seq analysis followed by RT-PCR and Sanger sequencing (Fig. S1A-S1C). The resultant transcript sequence was identical in the two cases at the fusion junction (exon 14 of *FOXP2* fused to exon 4 of *CPED1*), resulting in a stop-gain mutation right after the fusion breakpoint (Fig. S1C) and consequently yielding a predicted truncated FOXP2 protein with an aberrant C terminus. We subsequently characterized the genomic breakpoint of the *FOXP2*-*CPED1* in the fusion-positive samples using long-range PCR and whole-genome sequencing and revealed the mechanism as resulting from an ∼ 6.36-Mb fragment deletion occurred between *FOXP2* and *CPED1* on chromosome 7q3.1 (Fig. S1D-S1F).

We further observed that the exons of *FOXP2* before the fusion breakpoint (from 1st exon to 14th exon) were abundantly expressed in the fusion-positive sample (PC_1) determined by RNA-Seq (Fig. S1H), suggesting that most of the *FOXP2* expression took place from the fusion transcript rather than from natural *FOXP2*. Indeed, we validated very little natural *FOXP2* but the markedly increased *FOXP2* fusion transcripts, which resulted in a high-level of expression of truncated FOXP2 protein in the fusion-positive tumor (Fig. S1I-S1J)

To explore the potential mechanistic details underlying the increase in fusion transcript, we firstly performed reduced representation bisulfite sequencing (RRBS) analysis on the fusion positive tumor (PC_1) but did not observe low methylation status of *FOXP2* promoter region in this sample. Whole-genome sequencing analysis showed no somatic mutation in the *FOXP2* promoter of the tumor. Additionally, loss of the FOXP2 C-terminus seemed to have no effect on the stability of the protein [1].

Because the fusion led to the loss of the 3’ untranslated region (3’UTR) of *FOXP2* in the *FOXP2*-*CPED1* transcript (Fig. S1H-S1K), we thus asked whether the increased expression of the fusion gene could be due to an escape from targeted regulation of microRNA. We then conducted small RNA sequencing of PC_1 and combined our results with data from the TargetScan and MicroRNA databases. The overlap of the three datasets revealed 12 candidate microRNAs targeting *FOXP2*, but not *CPED1* (Fig. S1L). Luciferase reporter assays suggested that 2 of them, miR-27a and miR-27b, targeted *FOXP2* 3’UTR (Fig. S1M-S1N). miR-27a was recently revealed to target *FOXP2* involved in biological processes of gastric cancer [2]. Increased endogenous FOXP2 protein levels after transfection of miR-27a/b inhibitors in HEK 293T cells supported this suggestion (Fig. S1O). Furthermore, we added the *FOXP2* 3’UTR back into the *FOXP2*-*CPED1* construct and found decreased fusion protein expression upon transfection with miR-27a/b (Fig. S1P). All these data indicated that the absence of the 3’UTR of *FOXP2* in the fusion disrupted the regulation of microRNAs. Therefore, the absence of the 3’UTR of FOXP2 in the fusion disrupted the regulation of microRNAs, resulting in the increased FOXP2 fusion.

## Supplementary Figure Legends

**Figure S1 related to “Figure 1 - Figure supplement 1” and “Figure 1 - Figure supplement 2”.**
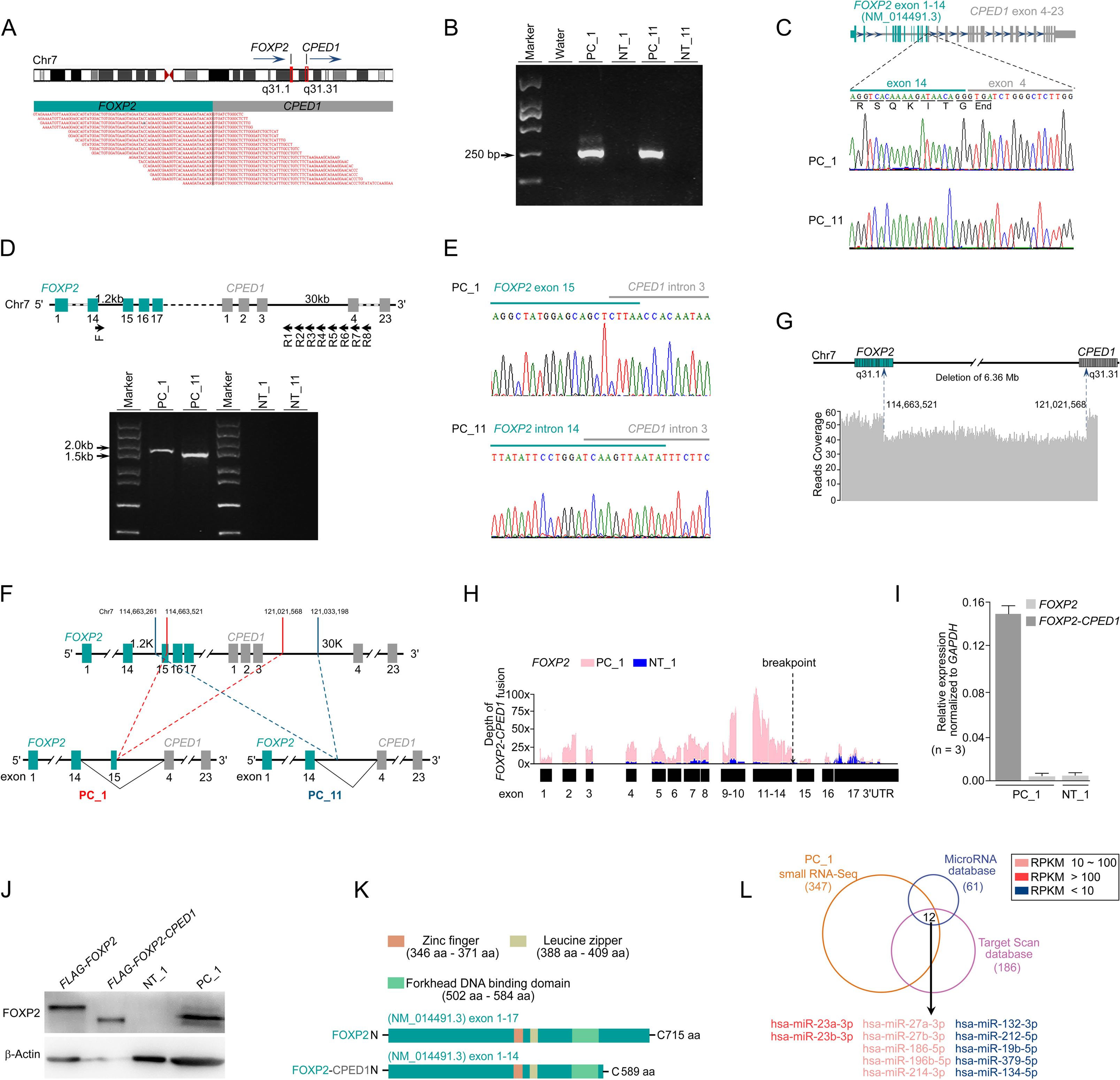
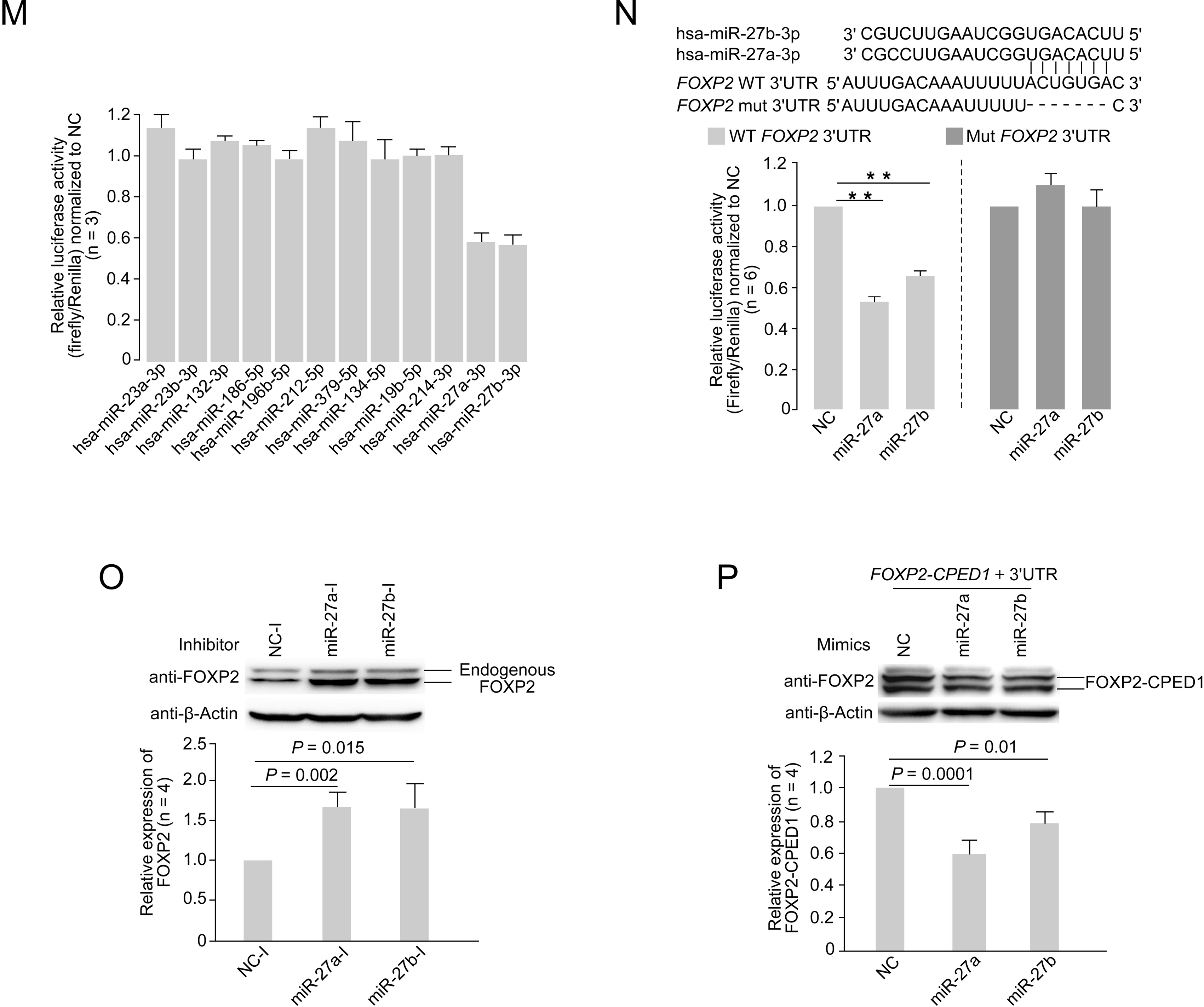
Identification of *FOXP2-CPED1* fusion gene in human prostate tumors. **A**. *FOXP2*-*CPED1* fusion gene identified in the prostate tumor of patient (PC_1) by RNA-Seq. *Top*, Schematic of chromosome 7 with the position and strand orientation indicated for the *FOXP2* and *CPED1* gene. *Bottom*, Schematic representation of the paired-end reads covering the junction between *FOXP2* and *CPED1*. **B-C**. **B**. *FOXP2*-*CPED1* specific PCR from cDNA derived from the prostate tumors (PC_1, PC_11) and their matched normal tissues (NT_1, NT_11), respectively. Marker, DNA marker. **C.** Sanger sequencing chromatogram of the PCR products as shown in (**B)** showing the reading frame encompassing the breakpoint in the two fusion-positive tumors (PC_1 and PC_11). **D.** *Upper panel*, A schematic of 8 primer sets (F-R1 to F-R8) for long-range PCR amplification for detection of the genomic breakpoint of *FOXP2*-*CPED1* fusion. *Lower panel*, The breakpoints of *FOXP2*-*CPED1* fusion at the genomic level by long-range PCR amplification of genomic DNA derived from the prostate tumors (PC_1, PC_11) and their matched normal tissues (NT_1, NT_11), respectively. Marker, DNA marker. **E.** Sanger sequencing chromatograms of long-range PCR amplification as shown in (**D)** shows the fusion breakpoints in genomic DNA from PC_1 or PC_11 that are positive for the *FOXP2*-*CPED1* fusion. **F.** A schematic of the genomic rearrangement pattern of the *FOXP2-CPED1* fusion in PC_1 and PC_11. Red dashed line, genomic fusion pattern of *FOXP2* exon 15 with intron 3 of *CPED1* in PC_1. Green dashed line, genomic fusion pattern of *FOXP2* intron 14 with intron 3 of *CPED1* in PC_11. **G.** Whole-genome sequencing of DNA from fusion-positive sample (PC_1) revealed an ∼ 6.36-Mb fragment deletion between *FOXP2* and *CPED1*. Dotted arrows indicate the breakpoints in the *FOXP2* and *CPED1* genes. **H.** Mapping of RNA-Seq reads to the *FOXP2* locus of a human reference genome (hg19) for the tumor sample (PC_1) and its matched normal tissue (NT_1). The dotted arrow indicates the fusion breakpoint in the *FOXP2* gene. **I**-**J**. Expression of natural *FOXP2* and *FOXP2*-*CPED1* fusion in the tumor (PC_1) and its matched normal tissue (NT_1) by qPCR (**I**) and by western blot analysis (**J**). Mean ± s.d. Overexpression of FLAG-*FOXP2*-*CPED1* or FLAG-*FOXP2* in NIH3T3 cells as positive controls. **K.** Schematic of protein structures of wild-type FOXP2 and truncated FOXP2 encoded by the *FOXP2*-*CPED1* fusion transcript. FOXP2-CPED1 fusion protein has an aberrant C-terminus but it retains the complete FOXP2 functional domains. **L.** The overlap of microRNAs identified in the *FOXP2*-*CPED1* fusion-positive sample (PC_1) by small RNA-Seq and predicted microRNAs targeting *FOXP2* from the TargetScan and MicroRNA databases. Total of 12 microRNA candidates are listed. **M.** Luciferase assay of the *FOXP2* 3’UTR after individual overexpression of 12 microRNA candidates in HEK 293T cells. Mean ± s.d.; n = 3. **N.** *Top*, Schematic indicating the location of the miR-27a and miR-27b binding sites in the wild-type *FOXP2* 3’UTR construct and the mutant with the binding site deleted. *Bottom*, Luciferase assay of the wild-type *FOXP2* 3’UTR versus mutant after transfection with miR-27a/b mimics or microRNA negative control (NC) in HEK 293T cells. \*\**P* < 0.0005 by 2-tailed Student’s *t* test, mean ± s.d.; n = 6. **O.** Detection of FOXP2 protein after transfection with the microRNA inhibitor negative control (NC-I), miR-27a inhibitor (miR-27a-I) or miR-27b inhibitor (miR-27b-I) in HEK 293T cells by western blotting. Histograms of quantitative analysis illustrating endogenous FOXP2 protein expression levels. *P* values calculated by 2-tailed Student’s *t* test, mean ± s.d.; n = 4. **P.** Detection of fusion protein after transfection of the *FOXP2*-*CPED1* plus *FOXP2* 3’UTR with NC, miR-27a or miR-27b in HEK 293T cells by western blotting. Histograms of quantitative analysis illustrating exogenous FOXP2-CPED1 truncated protein expression levels. *P* values calculated by 2-tailed Student’s *t* test, mean ± s.d.; n = 4.

**Figure S2 related to “Figure 1 - Figure supplement 3” and “Figure 1 - Figure supplement 4”.**
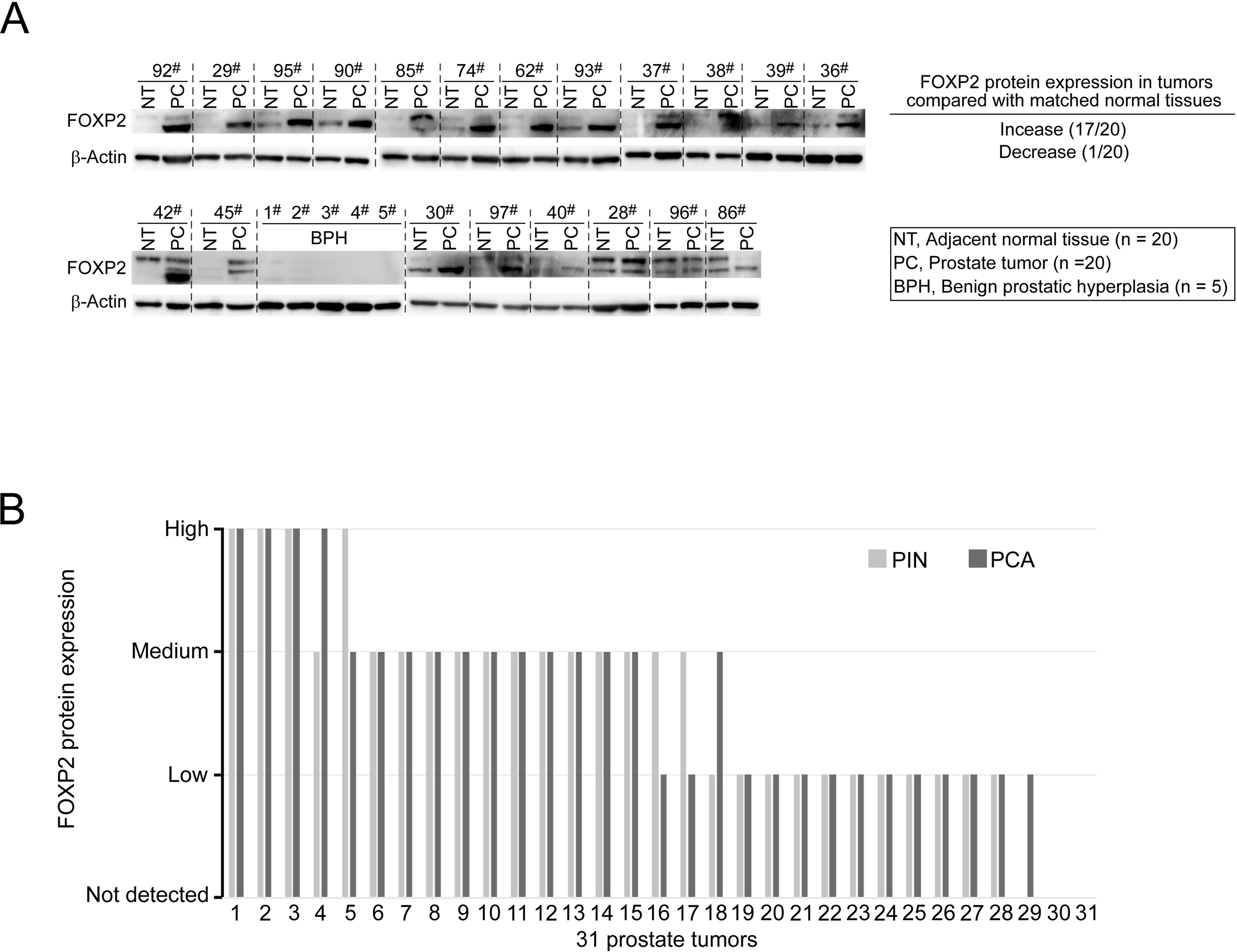
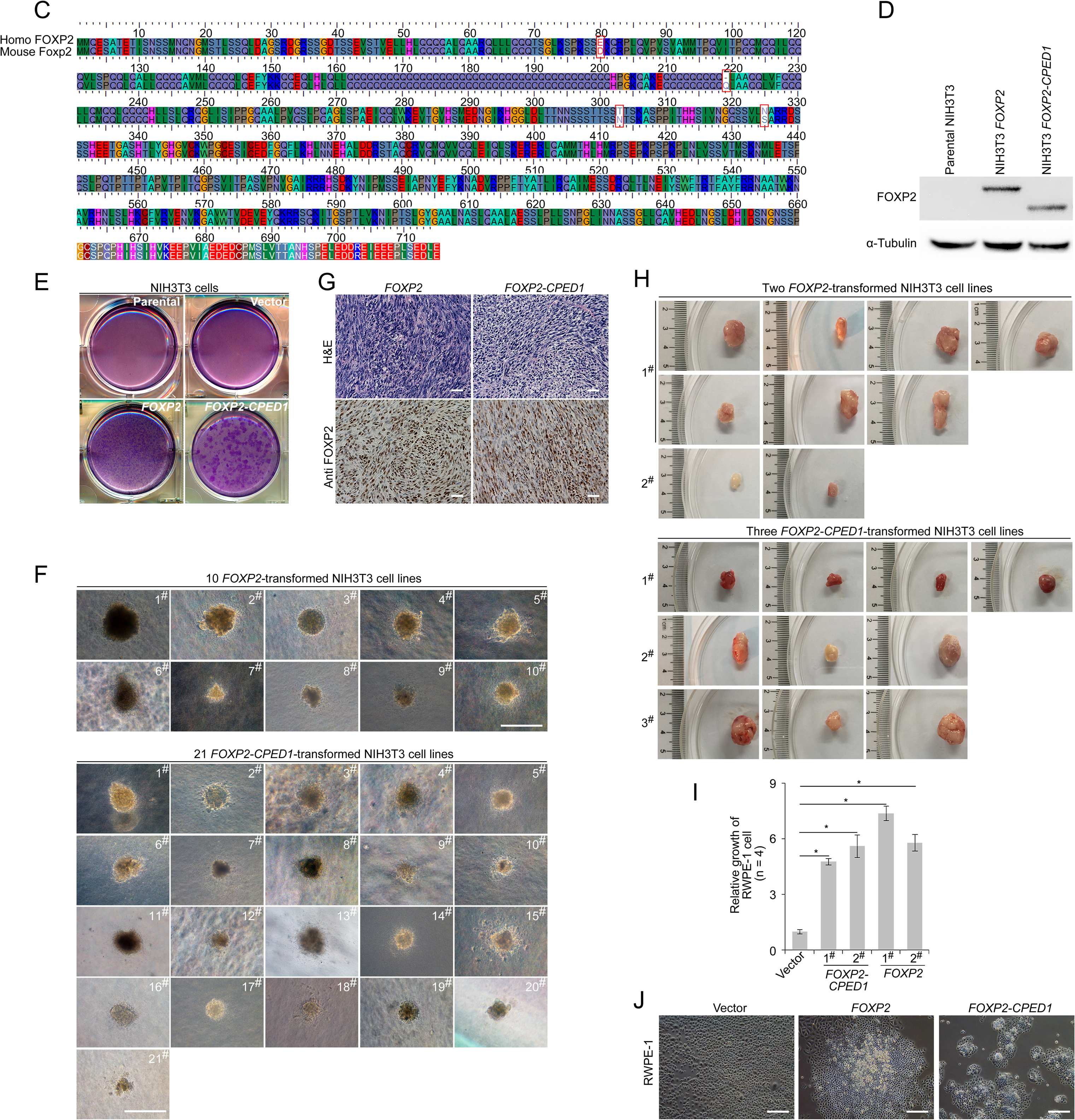
*FOXP2* functions as an oncogene in prostate cancer. **A.** Detection of FOXP2 protein expression levels in our primary prostatic adenocarcinoma (PCA) and the matched normal tissues (n = 20) and samples of benign prostatic hyperplasia (BPH) (n = 5) by western blotting. **B.** Evaluation of FOXP2 protein abundance in 31 prostate cancer specimens with the presence of prostatic intraepithelial neoplasia (PIN) adjacent to carcinoma by IHC. The 28 FOXP2- staining-positive cancers showed FOXP2 expression in the adjacent PIN as well. Two cases did not show FOXP2-staining in both cancers and the adjacent PIN. **C.** Alignment of homologs of human FOXP2 and mouse Foxp2 showing high conservation between the two species. **D.** Protein blot showing ectopic expression of FOXP2 and FOXP2-CPED1 fusion protein in NIH3T3 cells, respectively, and endogenic Foxp2 protein is undetectable in the parental NIH3T3 cells. **E.** Focus formation assay performed with parental NIH3T3 cells and NIH3T3 cells with stably expressing the empty vector, *FOXP2*-*CPED1* and *FOXP2*, respectively. **F.** All representative images of the soft agar colony formation assay showing transformed NIH3T3 cell lines induced by *FOXP2*-*CPED1* or *FOXP2* shown in Fig. 1F. Scale bars, 100 μm. **G.** Hematoxylin and eosin stain and immunohistochemical analysis using anti-FOXP2 antibody in tumors from NOD-SCID mice. Scale bars, 100 μm. **H.** Images showing tumors from mice injected with *FOXP2*-*CPED1* cell lines (1^#^, 2^#^ and 3^#^) and *FOXP2* cell lines (1^#^ and 2^#^) shown in Fig. 1H. **I.** Relative viability of cells with stably expressing *FOXP2* and *FOXP2-CPED1* (normalized to cells with stably expressing lentiviral empty vector) was determined using a Celltiter-Glo assay. RWPE-1 cells with stably expressing lentiviral empty vector, *FOXP2* and *FOXP2*-*CPED1*, respectively, were cultured for 6 days. \**P* < 0.005 by 2-tailed Mann-Whitney *U* test, mean±s.d.; n = 4. **J.** Focus formation assay performed with RWPE-1 cells with stably expressing lentiviral empty vector, *FOXP2* and *FOXP2*-*CPED1*, respectively. Scale bars, 100 μm.

**Figure S3 related to “Figure 2 - Figure supplement 1”.**
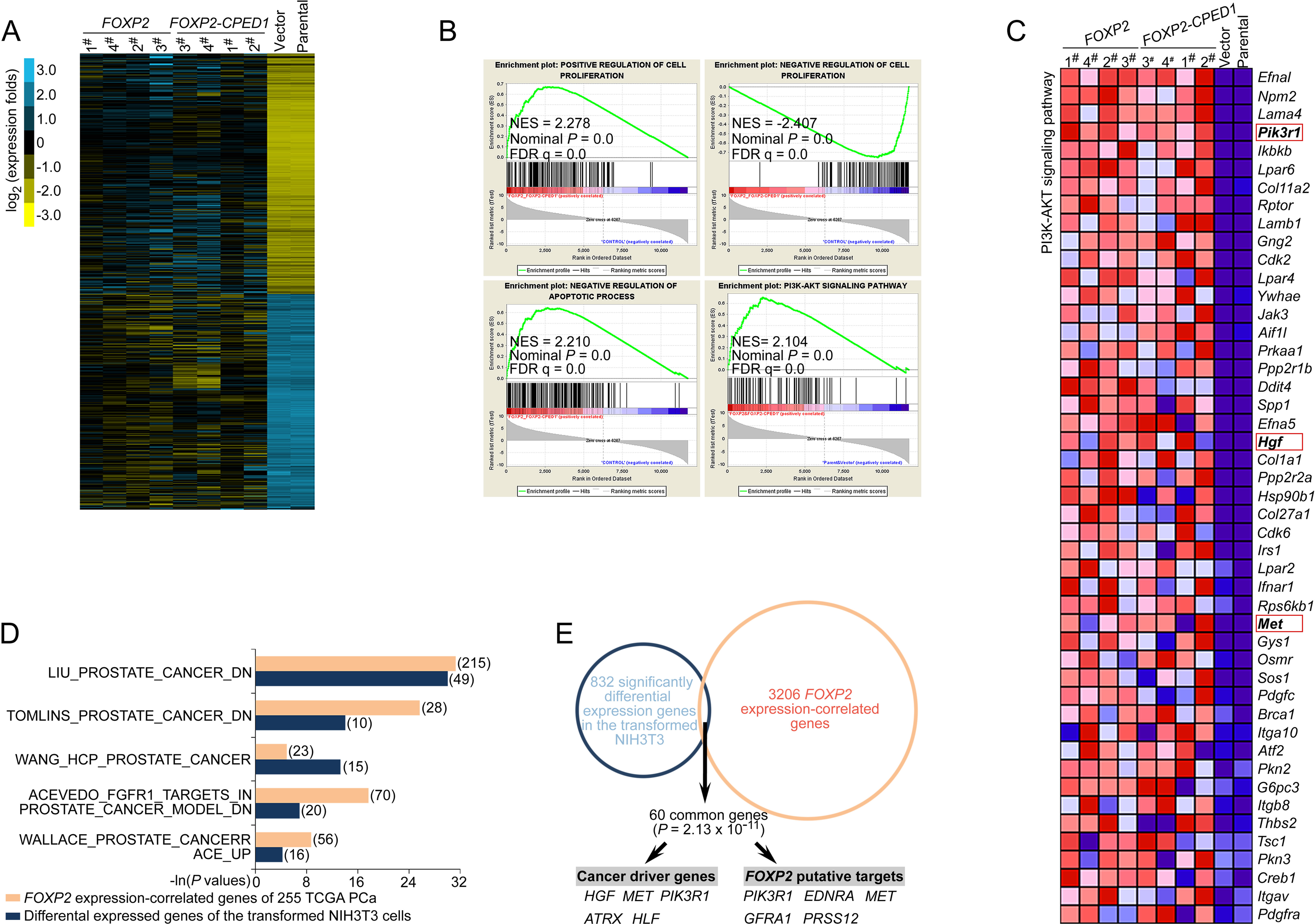
*FOXP2* activates oncogenic MET signalling in *FOXP2-*overexpressing Cells. **A.** Gene expression heat map of the significantly upregulated and downregulated genes from RNA sequencing analysis of NIH3T3 cells overexpressing *FOXP2* or *FOXP2-CPED1* relative to control cells. Yellow indicates high expression, blue indicates low expression. **B.** GSEA plots of the ranked list of differentially expressed genes in the *FOXP2*- or *FOXP2- CPED1*-transformed NIH3T3 cells and control cells (parental or stably expressing lentiviral vector NIH3T3 cells) were generated with four gene sets. NES, normalized enrichment score; FDR, false discovery rate. **C.** Heat map of differentially expressed genes contributing to the GSEA core enrichment of the PI3K-AKT pathway, involving Met signalling components (Pik3r1, Hgf and Met, marked by red rectangle) in the transformed NIH3T3 cells. Red, high expression; blue, low expression. **D.** GSEA analysis showing the enrichment of 832 differentially expressed genes in the *FOXP2* or *FOXP2-CPED1* transformed NIH3T3 cells and the enrichment of 3206 *FOXP2* expression-correlated genes (FECGs) in 255 human prostate cancer samples from the TCGA dataset (Prostate Adenocarcinoma, Provisional) in known prostate cancer gene sets from Molecular Signatures database. The number of enriched genes is indicated in parentheses. *P* values by 2-tailed Fisher’s exact test. **E.** The overlap between 832 significantly differentially expressed genes in the *FOXP2* or *FOXP2-CPED1*-transformed NIH3T3 cells and 3206 FECGs in human prostate tumors from the TCGA dataset (Prostate Adenocarcinoma, Provisional). Sixty common genes are represented in the overlap (*P* = 2.13 × 10^-11^ by 2-tailed Fisher’s exact test), including 5 cancer drivers and 5 *FOXP2* putative targets previously identified by ChIP-on-chip or ChIP assays.

**Figure S4 related to “Figure 4 - Figure supplement 1”.**
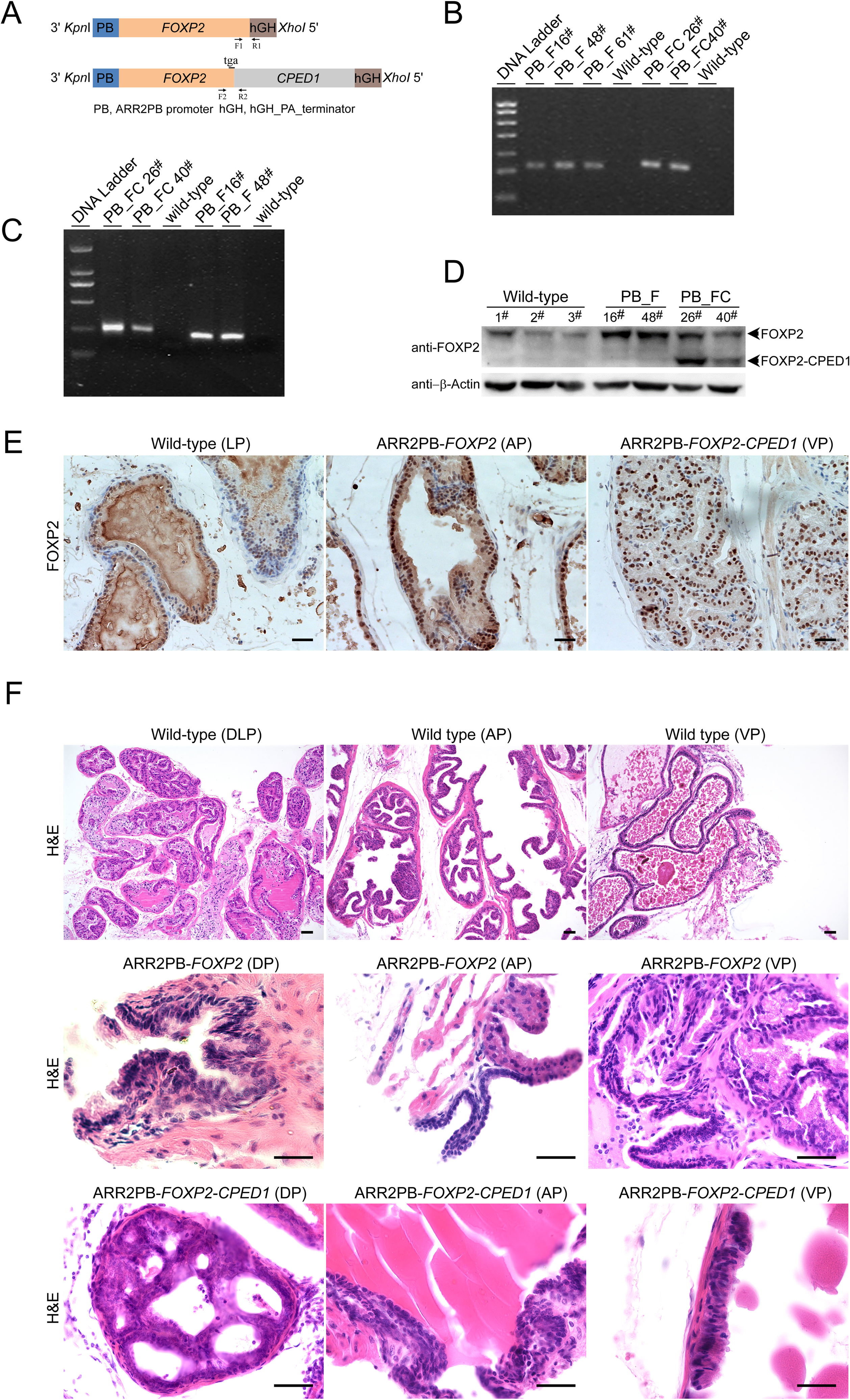
Generation of transgenic mice with prostate-specific expression of *FOXP2* or *FOPX2- CPED1*. **A.** Schematic of the *FOXP2* and *FOPX2-CPED1* transgene constructs with a composite ARR2PB promoter and a hGH_PA_terminator. Identification of ARR2PB*-FOXP2* or ARR2PB*-FOXP2-CPED1* transgenic mice at genomic and mRNA levels by PCR and RT-PCR using primers (F1 and R1, or F2 and R2). **B.** Genotyping of ARR2PB*-FOXP2* or ARR2PB*-FOXP2-CPED1* transgenic mice at genomic level using PCR. PB_F, ARR2PB*-FOXP2*; PB_FC, ARR2PB*-FOXP2-CPED1*. **C.** Specific detection of the mRNA expression levels of *FOXP2* or *FOXP2-CPED1* in prostates from both transgenic mice by RT-PCR. **D.** Detection of the protein abundance in prostates from transgenic mice and their littermate mice by immunoblot with an antibody against N terminus of FOXP2. **E.** Immunohistochemical staining for FOXP2 or truncated FOXP2 staining in the indicated prostate glands from ARR2PB-*FOPX2* and ARR2PB-*FOPX2-CPED1* transgenic mice (200×), demonstrating markedly increased protein expression in transgenic mice. AP, anterior prostate; VP, ventral prostate. **F.** Representative histological images of normal prostate glands in wild-type control mice (40×) and mPIN (200×) in ARR2PB-*FOPX2* and ARR2PB-*FOPX2-CPED1* transgenic mice at an average of 14 months of age. mPIN, murine prostatic intraepithelial neoplasia; AP, anterior prostate; VP, ventral prostate; DP, dorsalprostate; DLP, dorsal-lateral prostate.

**Figure S5 related to “Discussion - Figure supplement 1”.**
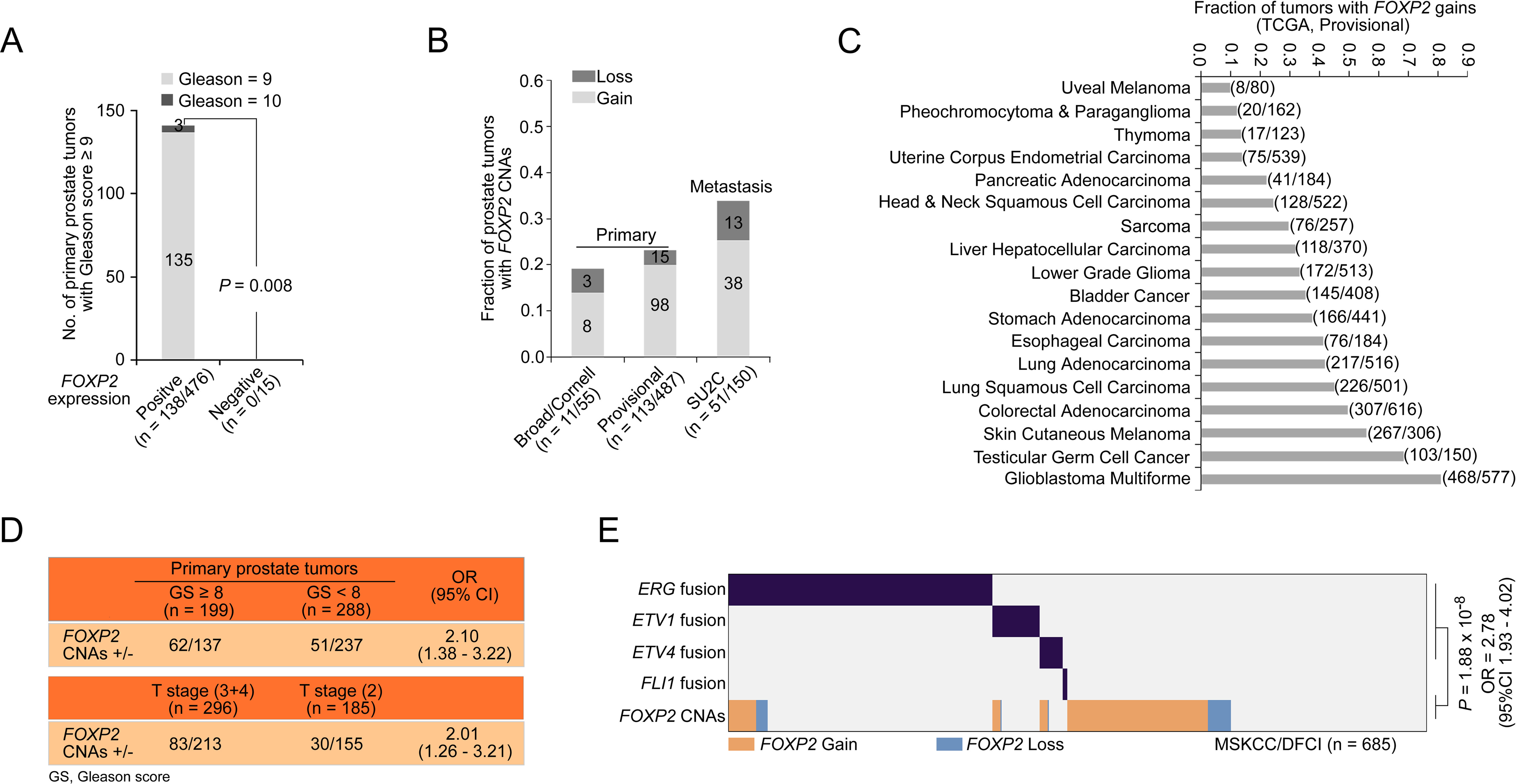
Clinical significance of *FOXP2* expression and its CNAs in prostate cancer. **A.** The relationship of *FOXP2* expression with Gleason score (≥ 9) in 491 primary prostate cancer samples from the TCGA dataset (Prostate Adenocarcinoma, Provisional) (*FOXP2* positive expression, n = 476 and *FOXP2* negative expression, n = 15). *P* value calculated by 2-tailed Fisher’s exact test. **B.** The fraction of tumors with *FOXP2* CNAs in two primary prostate cancer cohorts from the Broad/Cornell 2013 dataset (n = 55) and TCGA dataset (Prostate Adenocarcinoma, Provisional, n = 487) and in one metastatic tumors cohort from the SU2C dataset (n = 150). **C.** The fractions of *FOXP2* gains in 18 types of human solid tumors from the TCGA provisional datasets. The number of tumors with *FOXP2* gains versus total number of tumors is indicated in parentheses. **D.** The association of *FOXP2* CNAs with Gleason scores or T stages in primary prostate tumors from TCGA (Prostate Adenocarcinoma, Provisional, n = 487). *P* values were calculated using Fisher’s exact test. **E.** *FOXP2* CNAs exhibited a partially mutual exclusivity with ETS fusions in primary prostate tumors from the MSKCC/DFCI dataset (n = 685). *P* value calculated by 2-tailed Fisher’s exact test. **Source data** for Figure 1E, Figure 1F, Figure 3A, Figure 3B, Figure 3C, Figure 3F, Figure 3G, Figure S1J, Figure S1O, Figure S1P, Figure S2A, Figure S2D and Figure S4D.

